# Discovery and characterization of cross-reactive intrahepatic antibodies in severe alcoholic hepatitis

**DOI:** 10.1101/2023.02.23.529702

**Authors:** Ali Reza Ahmadi, Guang Song, Tianshun Gao, Jing Ma, Xiaomei Han, Mingwen Hu, Andrew M Cameron, Russell Wesson, Benjamin Philosophe, Shane Ottmann, Elizabeth A King, Ahmet Gurakar, Le Qi, Brandon Peiffer, James Burdick, Robert A Anders, Zhanxiang Zhou, Dechun Feng, Hongkun Lu, Chien-Sheng Chen, Jiang Qian, Bin Gao, Heng Zhu, Zhaoli Sun

**Affiliations:** Department of Surgery, Johns Hopkins University School of Medicine, Baltimore, United States; Department of Pharmacology and Molecular Sciences, Johns Hopkins University School of Medicine, Baltimore, United States; Department of Ophthalmology, Johns Hopkins University School of Medicine, Baltimore, United States; Laboratory of Liver Diseases, National Institute on Alcohol Abuse and Alcoholism (NIAAA), National Institutes of Health (NIH), Bethesda, United States; Department of Medicine, Johns Hopkins University School of Medicine, Baltimore, United States; Department of Pathology, Johns Hopkins University School of Medicine, Baltimore, United States; Center for Translational Biomedical Research and Department of Nutrition, University of North Carolina at Greensboro, North Carolina Research Campus, Kannapolis, United States; Department of Food Safety/Hygiene and Risk Management, National Cheng Kung University, Tainan city, Taiwan; Research Center, The Seventh Affiliated Hospital of Sun Yat-sen University, Shenzhen, P. R. China; Office of Translational Science, U.S. Food and Drug Administration, Silver Spring, United States

## Abstract

The pathogenesis of antibodies in severe alcoholic hepatitis (SAH) remains unknown. We sought to determine if there was antibody deposition in SAH livers and whether antibodies extracted from SAH livers were cross-reactive against both bacterial antigens and human proteins. We analyzed immunoglobulins (Ig) in explanted livers from SAH patients (n=45) undergoing liver transplantation and tissue from corresponding healthy donors (HD, n=10) and found massive deposition of IgG and IgA isotype antibodies associated with complement fragment C3d and C4d staining in ballooned hepatocytes in SAH livers. Ig extracted from SAH livers, but not patient serum exhibited hepatocyte killing efficacy in an antibody-dependent cell-mediated cytotoxicity (ADCC) assay. Employing human proteome arrays, we profiled the antibodies extracted from explanted SAH, alcoholic cirrhosis (AC), nonalcoholic steatohepatitis (NASH), primary biliary cholangitis (PBC), autoimmune hepatitis (AIH), hepatitis B virus (HBV), hepatitis C virus (HCV) and HD livers and found that antibodies of IgG and IgA isotypes were highly accumulated in SAH and recognized a unique set of human proteins as autoantigens. The use of an *E. coli* K12 proteome array revealed the presence of unique anti-*E. coli* antibodies in SAH, AC or PBC livers. Further, both Ig and *E. coli* captured Ig from SAH livers recognized common autoantigens enriched in several cellular components including cytosol and cytoplasm (IgG and IgA), nucleus, mitochondrion and focal adhesion (IgG). Except IgM from PBC livers, no common autoantigen was recognized by Ig and *E. coli* captured Ig from AC, HBV, HCV, NASH or AIH suggesting no cross-reacting anti-*E. coli* autoantibodies. The presence of cross-reacting anti-bacterial IgG and IgA autoantibodies in the liver may participate in the pathogenesis of SAH.

## Introduction

Severe alcoholic hepatitis (SAH) is a distinct clinical syndrome that can develop suddenly and quickly lead to liver failure. It carries a particularly poor prognosis with a 28-day mortality ranging from 30-50%^1,2,3^. Unfortunately, there is little to offer medically to such critically ill patients beyond supportive care with steroids, which improves survival in only a minority.

Studies have established connections between alcohol abuse, disruption of gut microbial homeostasis, and alcoholic liver disease (ALD). It has been speculated for over four decades that antibodies targeting intestinal microbes might play a role in pathogenesis of ALD^4,5,6,7,8,9^. For example, the presence of IgA and IgG on the cell membrane of hepatocytes was detected by direct immunofluorescence in patients with ALD, and the percentage of IgG-positive hepatocytes correlated with transaminase levels, independently of the histological findings^8^.

Although a number of studies demonstrated liver IgA deposition in ALD in the 1980s, other reports concluded that IgA deposition in the liver was not specific for ALD but might reflect the reduced metabolism of the damaged livers^10, 11^ or the clearance of excess IgA from the circulation^12^. A recent study^13^ confirmed that human livers contained IgA-secreting cells originating from Peyer’s patches and directed against intestinal antigens. Interestingly, livers from mice with ethanol-induced injury contain increased numbers of IgA-secreting cells and have IgA deposits in sinusoids^13^.

The primary aim of this study was to determine if there was antibody deposition in SAH livers and whether antibodies extracted from SAH livers exhibited hepatocyte killing efficacy. The second aim was to determine if antibodies deposited in the liver were cross-reactive antibodies against both bacterial antigens and human proteins and whether the cross-reactive antibodies were presented uniquely in SAH livers.

## Results

### Immunoglobulins in ballooned hepatocytes in SAH patients

To determine whether antibodies deposit in SAH livers, we collected explanted liver tissues from SAH patients during liver transplantation at Johns Hopkins. Liver tissue sections with H&E staining from SAH patients showed histologic features of SAH including macrovesicular steatosis, neutrophilic lobular inflammation, ballooning hepatocyte degeneration, Mallory-Denk bodies, and portal and pericellular fibrosis (***Figure 1A***). Immunohistochemistry (IHC) staining by using anti-human immunoglobulin antibodies demonstrated massive IgA and IgG deposition in ballooned hepatocytes in SAH livers, while none of the hepatocytes were stained with anti-human immunoglobulin antibodies in liver tissue sections from healthy donors except for positive staining in some hepatic sinusoid cells (***Figure 1B and C***). To further confirm the deposition of immunoglobulins in SAH livers, the presence of immunoglobulins in liver tissue homogenates form SAH (n=7) or healthy donors (n=7) was assessed by Western blot analysis and ELISA assays. Western blot analysis demonstrated that the levels of IgA and IgG were dramatically increased in all SAH livers as compared with the donor livers (***Figure 1D and E***). The IgM but not the IgE level was also significantly increased in SAH livers. The increase of IgA, IgG and IgM levels in SAH liver tissue homogenates was further confirmed by ELISA. IgA and IgG isotypes were major immunoglobulins in SAH livers (***Figure 1F***). Further analysis of IgG subclasses demonstrated that the IgG subclass levels - predominantly IgG1 – were significantly higher in SAH livers than that in healthy donors (***Figure 1G—figure supplement 1***). On the basis of these findings, we performed IHC staining for human IgG in SAH livers from 45 patients with liver transplantation and 10 donor livers in a clinical pathology lab at Johns Hopkins in a double-blind manner. The IgG+ hepatocytes in scanned slides of stained tissues sections were analyzed by using HALO^TM^ Image Analysis Software. Positive cells were reported as percentage stained surface area of total annotated area by digital analysis (***figure supplement 1***). Few IgG+ hepatocytes were identified in donor livers. In contrast, on average ∼40% of the hepatocytes (ranging from 4% to 80%) were IgG positive in the SAH livers (***Figure 1H***). These findings demonstrated the deposition of immunoglobulin antibodies in ballooned hepatocytes in SAH livers.

**Figure 1.**
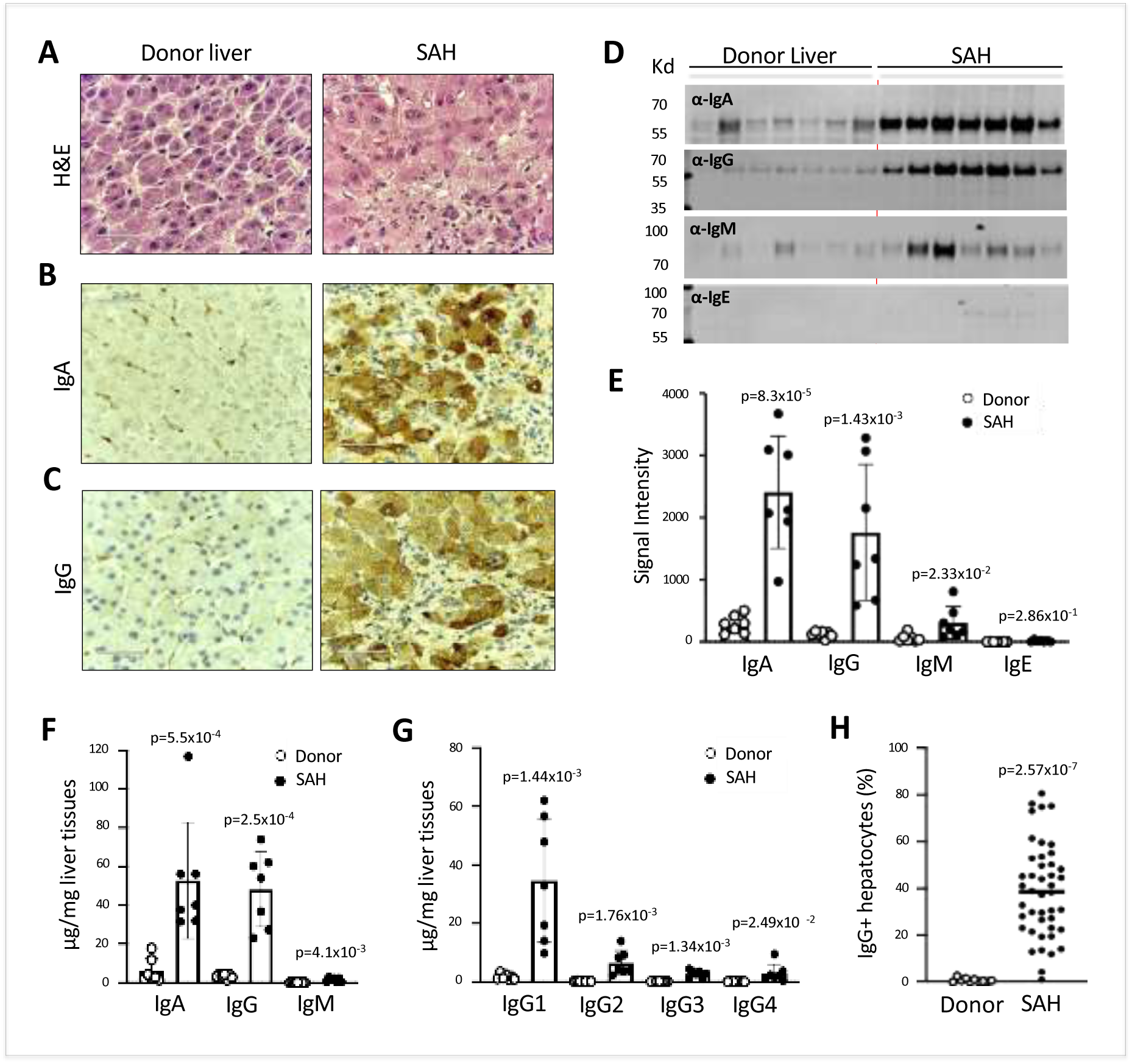
Immunoglobulin (Ig) deposition in ballooned hepatocytes of explanted livers from SAH patients. (A) Liver tissue sections with H&E staining showed histologic features of SAH. b,c, Immunohistochemistry staining by using anti-human IgA (B) or IgG (C) antibodies demonstrated IgA and IgG deposition in ballooned hepatocytes in SAH livers. Representative tissue sections from 45 SAH or 10 healthy donor (HD) livers. (D-E) Immunoglobulin levels in liver tissue homogenates from SAH or HD (n=7/group) were quantified by Western blot analysis (D). Western blot analysis demonstrated that the levels of IgA, IgG and IgM were significantly increased in SAH livers as compared with the HD livers (E). (F-G) Immunoglobulin isotypes (f) and IgG subclass levels (G) were quantified by ELISA (n=7/group). (H) IgG positive hepatocytes in tissues sections from 45 SAH patients and 10 healthy donors were quantified by immunohistochemistry staining and using HALO^TM^ Image Analysis Software. Figure supplement 1. Immunoglobulin deposition in alcoholic cirrhotic livers.

### Immunoglobulin deposition is associated with activation of complement in ballooned hepatocytes and immunoglobulin extracted from SAH livers exhibits hepatocyte killing efficacy *in vitro*

IgG, especially IgG1, plays a critical role in the classical completement activation pathway. To determine if immunoglobulin in ballooning hepatocytes induces activation of complement, C3d and C4d were analyzed in SAH livers. IHC staining showed the presence of both C3d and C4d in ballooning hepatocytes in SAH livers but not in the donor livers (***Figure 2A and 2B***). Double staining for IgG and complement fragments C3d or C4d showed IgG co-stained with C3d or C4d in ballooning hepatocytes (***Figure 2C and 2D***). These results indicated that IgG deposition in hepatocytes was associated with activation of complement. Furthermore, complement activation including the presence of C3d and C4d in SAH liver was confirmed by Western blot analysis (***Figure 2E and 2F***). Finally, we asked whether antibodies (Ig) extracted from SAH livers exhibit hepatocyte killing efficacy in an ADCC assay. Compared to isotype control human Ig from healthy donors, no increased hepatocyte killing was observed when PBMCs (effector cells) from healthy donors were added into cultured human hepatocytes (target cells) in the presence of serum Ig from SAH patients. However, the hepatocyte killing efficacy was significantly increased when the same levels of Ig extracted from SAH livers were added into the hepatocytes/PBMCs co-culture system (***Figure 2G***). These results demonstrated that immunoglobulin antibodies deposited in hepatocytes of SAH livers could induce activation of complement but more importantly, exhibited antibody-dependent cellular cytotoxicity of hepatocytes. Therefore, deposition of immunoglobulin antibodies may contribute to the hepatocyte ballooning degeneration and necrotic damage in SAH.

**Figure 2.**
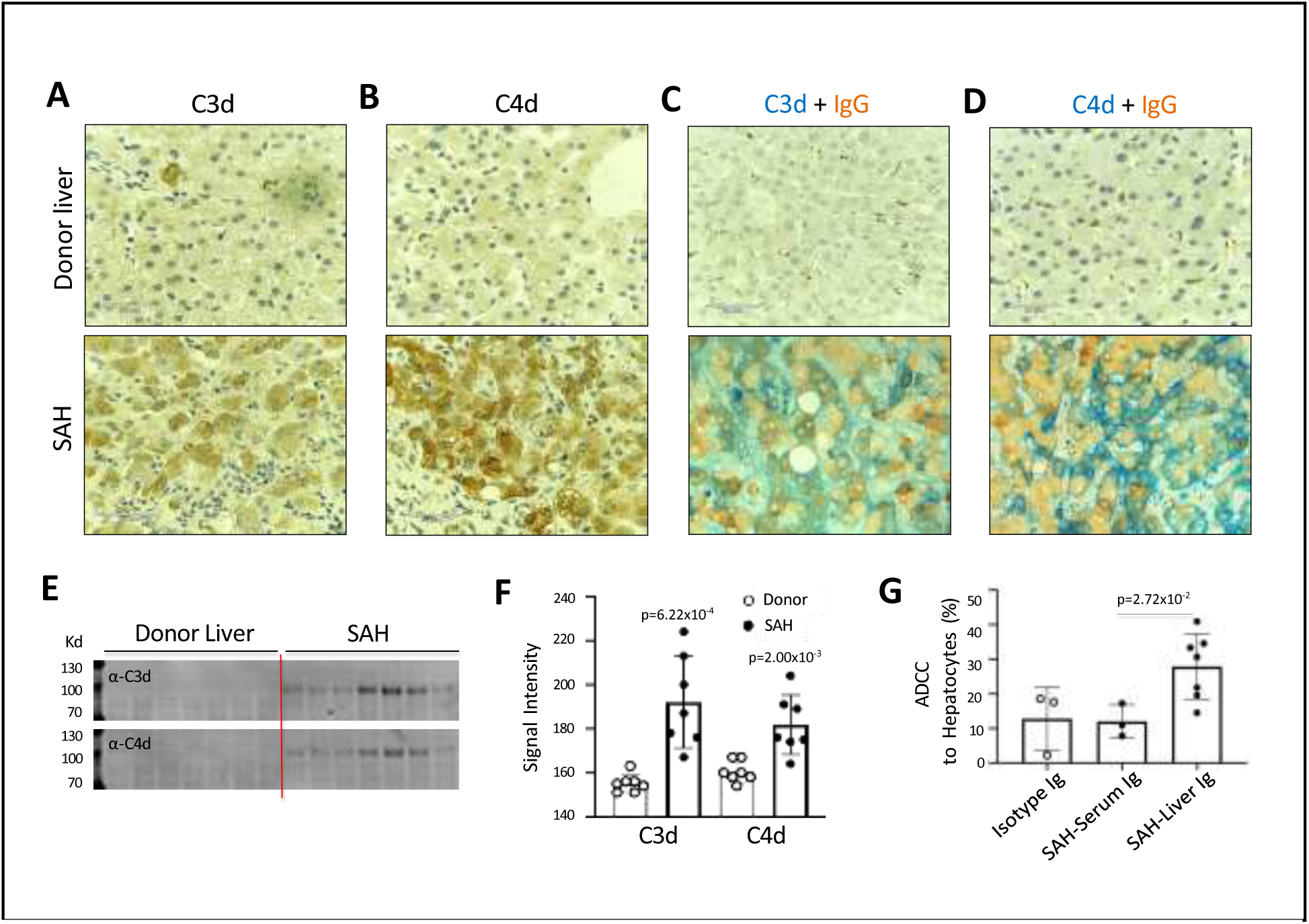
Ig deposition is associated with activation of complement in ballooned hepatocytes and Ig extracted from SAH livers exhibits hepatocyte killing efficacy *in vitro.* To determine if immunoglobulin in ballooning hepatocytes induces activation of complement, complement fragments C3d and C4d were analyzed in SAH livers. (**A-B**) Immunohistochemistry staining showed the presence of both C3d (**A**) and C4d (**B**) in ballooning hepatocytes in SAH livers but not in the donor livers. (**C**) Double staining for IgG and complement fragments C3d or C4d showed IgG co-stained with C3d or C4d in ballooning hepatocytes. Representative tissue sections from 7 samples per group. (**E-F**) C3d and C4d levels in SAH livers were quantified by Western blot analysis (n=7). **g**, Ig extracted from SAH livers but not serum exhibit hepatocyte killing efficacy in ADCC assay. Representative data from three independent experiments.

To further define the Ig and C4d deposition on the membrane of ballooned hepatocytes, we performed multiplex analyses by co-staining liver tissue sections with multiple cell markers for hepatocytes (Heppar1), Kupffer cells (CD68), hepatic sinusoid endothelial cells (CD32) and hepatocyte membrane (β-catenin). Images of confocal microscopy showed the presence of both IgG and IgA in hepatic sinusoid endothelial cells but not hepatocytes in the donor livers, while the majority of ballooned hepatocytes co-stained with IgG and IgA in SAH livers (new ***Figure 3A***). Further, co-staining with β-catenin detected heavy IgG and IgA deposition on hepatocyte membrane (***Figure 3B***). Interestingly, the IgG and IgA on hepatocyte membrane were co-stained with C4d as well (***Figure 3C***).

**Figure 3.**
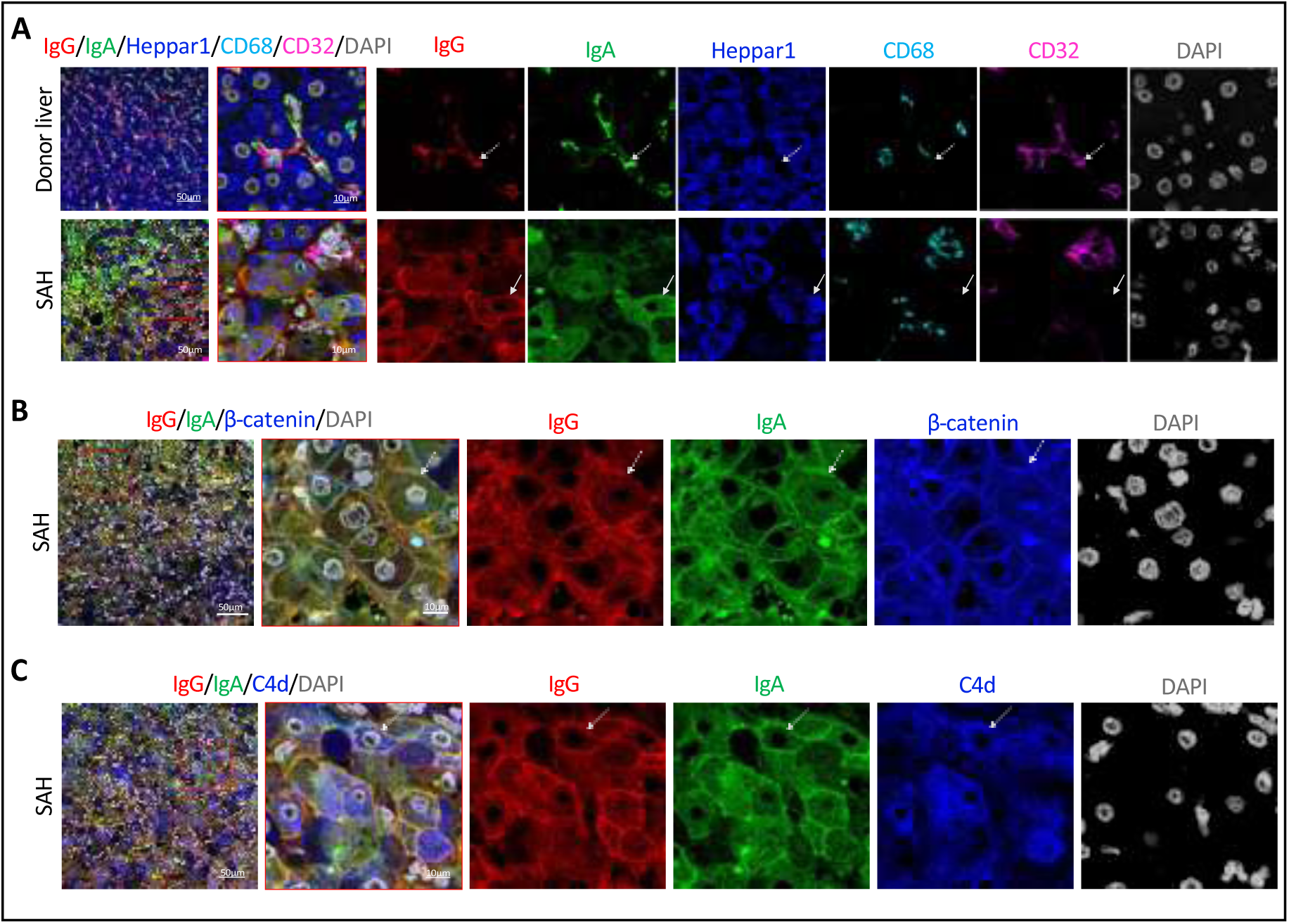
To determine if Ig recognize cell surface antigens of ballooned hepatocytes, multiplex immunofluorescence staining was performed in liver tissue sections. (**A**) Images of confocal microscopy showed the presence of both IgG (red) and IgA (green) in ballooning hepatocytes (blue) in SAH livers (lower panels), while only hepatic sinusoid endothelial cells (CD32+, purple) stained with IgG and IgA in donor livers (upper panels). (**B**) Co-staining with β-catenin (blue) demonstrated IgG (red) and IgA (green) deposition on membrane of ballooning hepatocytes in SAH livers. (**C**) Triple staining for IgG (red), IgA (green) and C4d (blue) showed both IgG and IgA co-stained with C4d on the surface of hepatocyte. Representative tissue sections from 6 samples per group.

### Human proteome array-identified autoantigens were recognized by immunoglobulins extracted from SAH livers

We used the human proteome microarray (HuProt), comprised of 21,240 individual purified human proteins, to perform antibody profiling assays^15^. Each liver specimen was treated to release tissue-deposited immunoglobulins (***Figure 4A***). After neutralization, the extracted antibodies from each liver sample were separately probed to the HuProt arrays, using isotype-specific secondary antibodies to obtain the IgG, IgA, IgM, and IgE autoimmune signatures of the same liver sample (***Figure 4B***). Many positive human proteins were recognized by each of the four Ig isotypes in all five SAH samples. Each antibody profiling assay was performed in duplicate and only those reproducible signals were scored. A substantial fraction of autoantigens was shared by the antibodies in all five SAH livers, regardless of the Ig isotype (***Figure 4C—figure supplement 2***). The total numbers of the shared autoantigens recognized by the IgG, IgA, IgM, and IgE isotypes were 346, 319, 194, and 10 (Suppl. Figure S2), respectively, suggesting that the shared autoantibodies of the IgG and IgA isotypes were the most prevalent, while the counterparts of IgE isotype were the scarcest. With tissues of five donor livers, the numbers of positive human proteins were much lower (***Figure 4C—figure supplement 3***).

**Figure 4.**
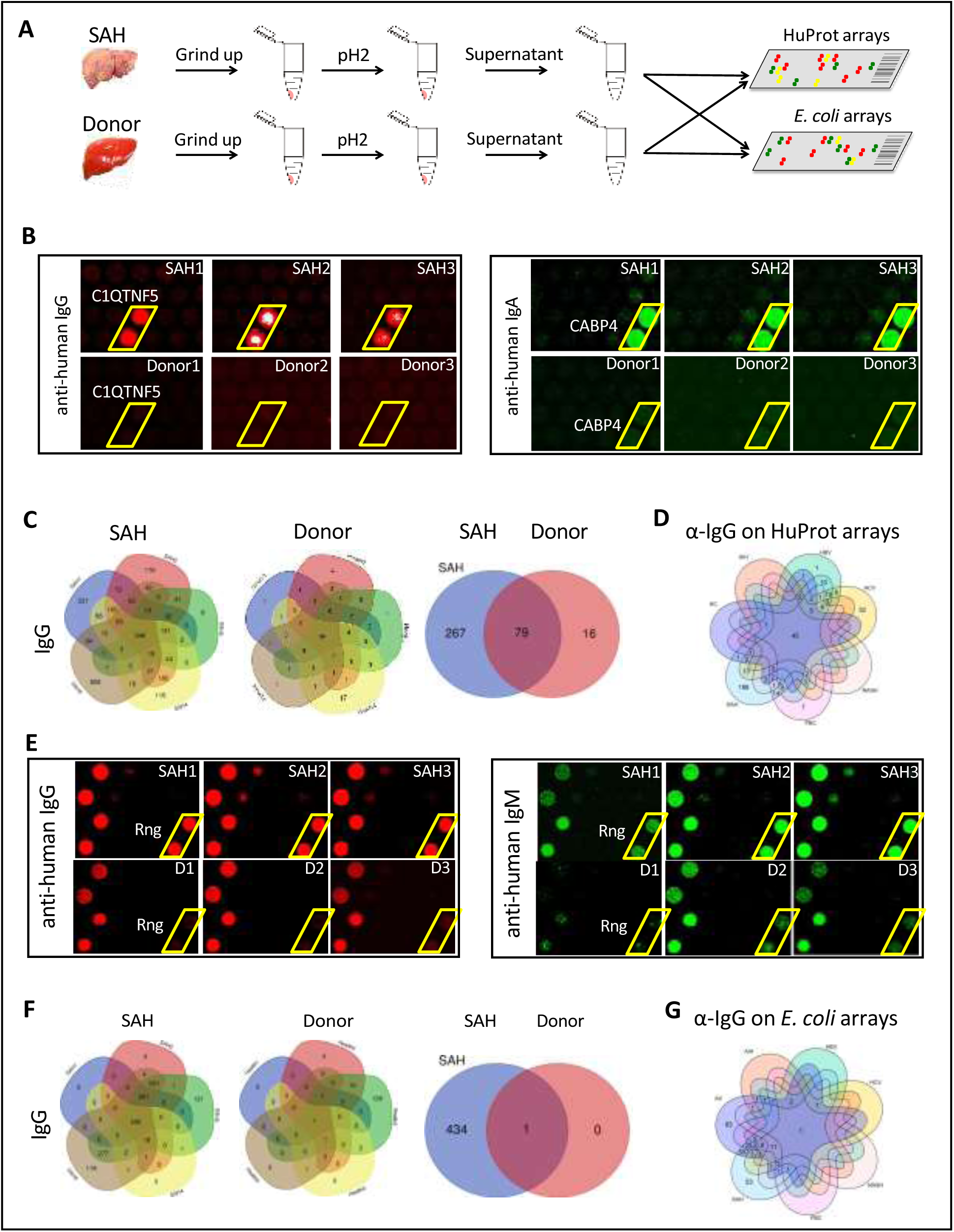
Human and *E. coli* proteome arrays identify a group of unique antibodies in the SAH liver recognize both human and bacterial antigens. (A) Each liver tissue piece was ground up and treated under low pH to release tissue-deposited Ig. After neutralization, the extracted Ig from each liver sample were separately probed to the HuProt or E. coli protein arrays, followed by incubation with the isotype-specific secondary antibodies to obtain the Ig isotype immune signatures of the same liver sample. (B) Representative images of HuProt arrays. (C) Venn diagram analysis to identify shared autoantigens of each liver disease. 346 autoantigens were shared by the IgG antibodies in all five SAH livers (left panel), 95 autoantigens were shared by the IgG antibodies in all five HD livers (middle panel), and a large fraction (i.e., 267 proteins) of the SAH-shared IgG autoantigens was not found in the HD livers (right panel), suggesting existence of a SAH-specific autoimmune signature. (D) A seven-way Venn diagram analysis showed that 45 autoantigens were commonly recognized by IgG isotype autoantibodies in liver tissue homogenates extracted from different liver diseases, while 188 unique IgG autoantigens were recognized by tissue homogenates from the SAH livers. (E) Representative images of *E. coli* protein arrays. (F) Venn diagram analysis to identify shared *E. coli* antigens recognized by each liver disease. 435 *E. coli* antigens were commonly recognized by IgG antibodies in the five SAH livers (left panel), while only 1 *E. coli* antigen was commonly recognized by IgG antibodies in the five HD livers (middle panel). 434 out of 435 *E. coli* antigens were uniquely recognized by IgG antibodies in SAH livers but not HD livers (right panel). (G) A seven-way Venn diagram analysis showed that unique IgG bacterial antigens were only identified by using liver tissue homogenates from SAH or AC. Figure supplement 2. Numbers of autoantibodies shared by all five SAH livers. Figure supplement 3. Venn diagram analysis of all liver samples. Figure supplement 4. Unique autoantigens recognized by Ig from diseased livers on the HuProt arrays. Figure supplement 5. Venn diagram analysis of antigens identified on the *E. coli* proteome arrays. Figure supplement 6. Unique bacterial antigens recognized by Ig from diseased livers on the *E. coli* proteome arrays.

Although 95 shared IgG autoantigens were identified by the donor liver samples, 79 (83.2 %) of them were also shared by the SAH livers (***Figure 4C***). More importantly, a large fraction (i.e., 267 proteins) of the SAH-shared autoantigens were not found in the donor livers (***Figure 4C***)

We applied the above approach to a group of five livers explanted from patients with AC, AIH, PBC, NASH, HCV, and HBV-infection. SAH still exhibited the highest number of shared IgG autoantigens, while HBV showed highest numbers of shared IgA and IgE autoantigens, and PBC showed the highest number of shared IgM autoantigens (***figure supplement 3***). The numbers of shared IgG, IgA and IgM autoantigens were much higher in SAH livers (n=859) as compared with AC (n=349), HBV (n=735), other liver diseases (n<428) or HD livers (n=448). Although the shared IgE autoantigens were the lowest in all seven liver diseases, each disease showed a distinct autoantibody signature (***figure supplement 3***).

We compared the shared autoantigens recognized by SAH immunoglobulins to their counterparts from the other six liver diseases. By Venn diagram analysis 45, 41, 68 and 8 autoantigens were commonly recognized by IgG, IgA, IgM and IgE isotype autoantibodies in liver tissue extracted from different liver diseases (***figure supplement 4***). 188 unique IgG autoantigens, 45 unique IgA autoantigens and 7 unique IgM autoantigens were recognized by tissue homogenates from the SAH livers, whereas the second highest in this category was HBV livers in which 1 unique IgG autoantigen, 88 unique IgA autoantigens and 25 unique IgM autoantigens were identified (***Figure 4D— figure supplement 4***). The third in this category was PBC in which 7 unique IgG autoantigens, 2 unique IgA autoantigens and 81 unique IgM autoantigens were identified. The tissue homogenates from HCV livers recognized 32 unique IgG autoantigens, 4 unique IgA autoantigens and 3 unique IgE autoantigens, while only 4 unique IgA autoantigens were recognized by tissue homogenates from AC livers (***figure supplement 4***), showing disease-distinct autoantibodies in diseased livers regardless of their etiology. A large number of unique autoantigens were recognized by Ig recovered from SAH livers (***table supplement 1***) indicating immunoglobulins (especially IgG) deposited to the SAH livers (***Figure 1***) might play an important role in pathogenesis.

### Immunoglobulins from SAH or AC livers recognize a unique set of bacterial antigens

Antibodies secreted into the gut mostly target bacteria and bacterial products^21^. Immunoglobulins deposited in SAH livers might be from the gut, and these immunoglobulins may be antibodies targeting intestinal bacterial antigens. To test this, we employed a bacterial proteome array, comprised of 4,256 purified *E. coli* proteins encoded by a commensal strain K12, to do antibody profiling assays^16^ (***Figure 4E***). Using the same liver tissues and approach described above, we obtained the immune signatures of the 40 livers from 7 diseases and 5 HD in duplicate.

Many more bacterial than human antigens were recognized by immunoglobulins from alcoholic livers (AC and SAH) compared with healthy donor livers (***figure supplement 5***). The differences in shared bacterial antigens between alcoholic livers and healthy donor livers were more pronounced across all four Ig isotypes. For instance, only one bacterial antigen was commonly recognized by IgG antibodies from HD livers, but 435 shared bacterial antigens (about 10.2% of 4,256 purified *E. coli* proteins) were recognized by IgG antibodies in the five SAH livers (***Figure 4F***), and 466 shared bacterial antigens were recognized by IgG antibodies in the five AC livers (***figure supplement 5***). The numbers of the commonly shared bacterial antigens identified by the IgG antibodies extracted from the other five liver diseases were much lower, ranging from 7 to 63 (***figure supplement 5***). In addition, the numbers of IgA, IgM or IgE recognized bacterial antigens by liver tissue homogenates from SAH or AC were much higher than that in other liver diseases, except PBC showing a higher number of IgM antigens than AC (***figure supplement 5***). This observation suggests that chronic or acute alcoholic liver disease is associated with specific antibacterial antibody signatures.

A large number of shared bacterial antigens were recognized by liver tissue homogenates from each liver disease. We found that 1, 6, 24 and 5 *E. coli* antigens were commonly recognized by IgG, IgA, IgM and IgE isotype antibodies in liver tissue homogenates extracted from different liver diseases (***Figure 4F***—***figure supplement 6***). Unique IgG or IgA bacterial antigens were only identified by using liver tissue from SAH or AC (***Figure 4G***—***figure supplement 6***). More unique bacterial antigens were recognized by the SAH than AC IgA (110 vs. 54), IgM (54 vs. 1) and IgE (45 vs. 1), but less so by the SAH IgG (53 vs. 83) (***table supplement 2***). Notably, 69 unique IgM bacterial antigens were recognized by PBC liver tissue. Common anti-*E. coli* protein antibodies are present in diseased livers regardless of etiology, but unique anti-*E. coli* IgG and IgA antibodies exist only in alcoholic livers predominantly in SAH.

### Immunoglobulins with anti-*E. coli* antigen specificities in SAH livers recognize human antigens

The prevalent anti-bacterial immunoactivity of the immunoglobulins in SAH livers suggested that immunoglobulins deposited in SAH livers might be derived from leaky gut and these anti- bacterial antibodies cross-react with human liver proteins. We used proteins from *E. coli* (strain K12) immobilized on magnetic beads to capture immunoglobulins pooled from the SAH livers (***Figure 5A***) (*E. coli* captured-Ig) which were then released and incubated on the HuProt arrays to determine whether these bacterium-recognizing antibodies could also cross-react with human proteins (examples shown in ***Figure 5B***). At a stringent cutoff value (e.g., S.D.=10) 694, 796, 451 and 42 human proteins were reproducibly identified by E. coli protein captured IgG, IgA, IgM and IgE antibodies from the five SAH livers, respectively (***Figure 5C*** and antibody recognized protein sets in ***table supplement 3***). Strikingly, many of these E. coli binding Ig recognized proteins (74.06%, 76.46%, 48.39% and 49.41% respectively) were also found to be the autoantigens recognized by immunoglobulins recovered directly from the five SAH livers.

**Figure 5.**
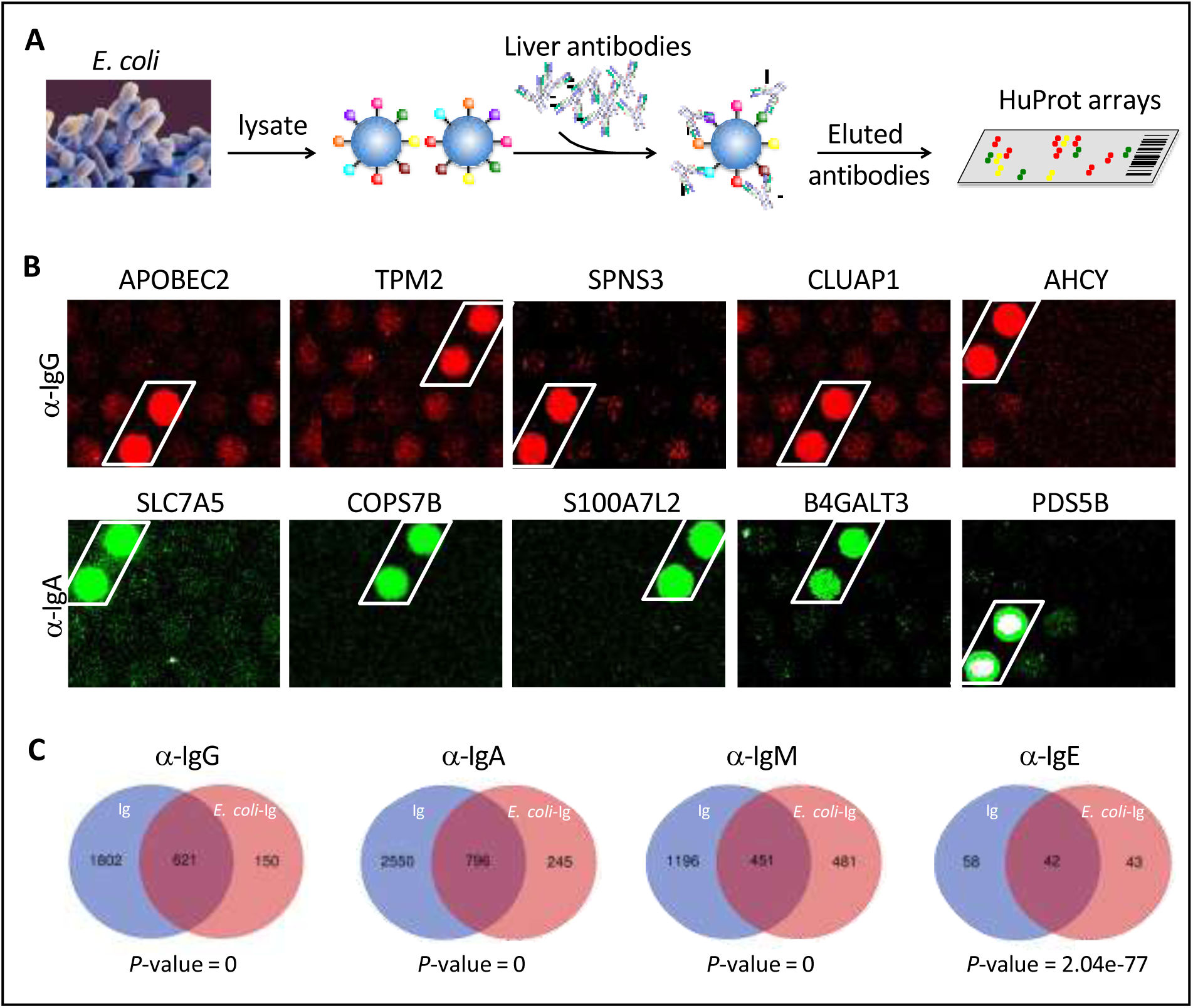
*E. coli* antigens-captured Immunoglobulins from SAH livers recognize human protein antigens. (**A**) To determine if anti-bacterial antibodies cross-react with human proteins in the liver, total proteins from *E. coli* (strain K12) were extracted and immobilized on NHS-activated magnetic beads to capture immunoglobulins pooled from the five SAH livers. *E. coli* protein-captured antibodies (*E. coli*-Ig) were then released and incubated on the HuProt arrays. (**B**) Representative images of *E. coli*-Ig on HuProt arrays. (**C**) 771, 1041, 932 and 85 human proteins were reproducibly identified by *E. coli* protein captured IgG, IgA, IgM and IgE antibodies from the five SAH livers. Venn diagram analysis showed many of these proteins (621/771, 796/1041, 451/932 and 42/95 respectively) were also found to be the autoantigens recognized by immunoglobulins recovered directly from the five SAH livers.

These results demonstrated that there exist a large number of cross-reacting antibodies that recognize both bacterial and human proteins in the SAH livers.

### Gene ontology (GO) enrichment analysis of proteome arrays identified autoantigen enriched unique common cellular components recognized by both Ig and *E. coli*-captured Ig in SAH livers

To determine if autoantigens recognized by Ig from diseased livers were specifically presented in cellular components, we performed GO enrichment analysis on antibody recognized protein sets. Proteins recognized by IgG antibodies from SAH livers were significantly over-represented in 8 cellular components including cytosol, cytoplasm, nucleus, mitochondrion, focal adhesion extracellular exosome, mitochondrial intermembrane space and ruffle membrane, while autoantigens recognized by IgA antibodies from SAH livers were over-represented in cytosol and cytoplasm (***Figure 6A***—***table supplement 5***). These proteins are involved in a number of biological processes including nucleobase-containing small molecule interconversion, protein phosphorylation, signal transduction, regulation of cell proliferation, mitochondrial ribosome reassembly and mitochondrion organization (***Figure 6B and C***). With the exception that cellular components were identified by IgM antibodies from PBC livers (***table supplement 5***), no cellular component was recognized by Ig from other diseased livers. Interestingly, *E. coli*-captured IgG and IgA from SAH livers recognized 5 out of 8 cellular components which were recognized by Ig directly extracted from SAH livers (***Figure 6D***). This was observed only in SAH livers (***table supplement 4***). These results further demonstrated the presence of unique cross-reacting anti-*E. coli* autoantibodies in the SAH livers.

**Figure 6.**
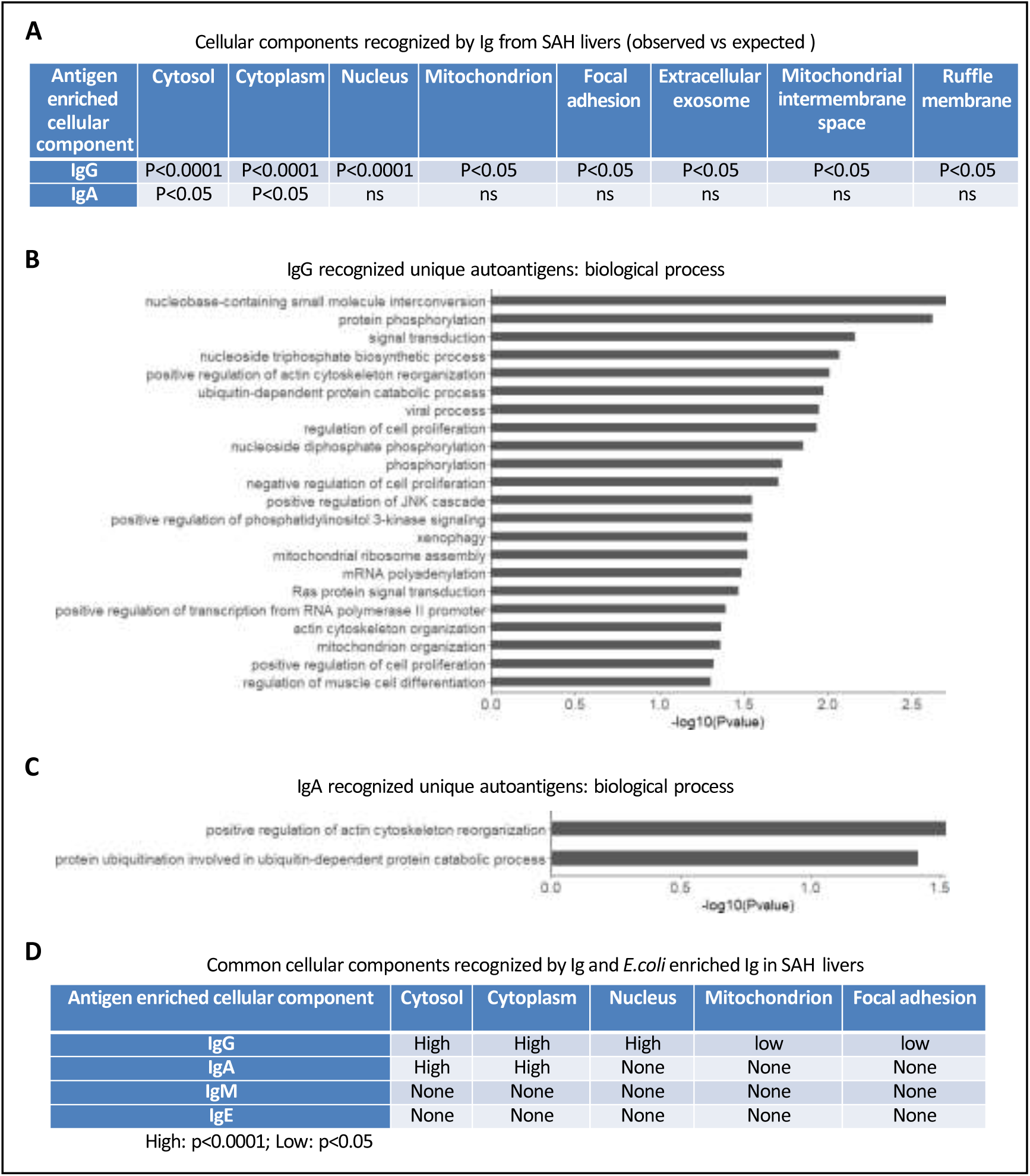
Gene ontology (GO) enrichment analysis of proteome arrays identifies autoantigen enriched common cellular components recognized by both Ig and E. coli captured Ig in SAH livers. (**A**) Cellular components recognized by IgG and IgA antibodies in SAH livers. (**B-C**) Biological processes are involved by IgG autoantigens (**B**) and IgA autoantigens (**C**). (**D**) Common cellular components recognized by both Ig and E. coli antigens captured Ig in SAH livers.

### The infiltration of B and plasma cells in SAH livers is associated with increased immunoglobulin gene expression

Alcohol-derived leaky gut may promote translocation of gut bacterial products and Peyer’s patches IgA-secreting plasma cells to the liver^13^. To determine if the migration of bacteria and/or bacterial products from the bowel to liver occurred in SAH, we performed immunohistochemistry staining for the gram-negative bacterial (*E. coli*) product livers. Both LPS and LTA were in liver tissue from SAH patients cf. controls (***Figure 7A***), especially in the inflammatory areas. The increase of LPS levels in SAH liver tissues was confirmed by using Pierce™ Chromogenic Endotoxin Quant Kit (***Figure 7B***).

**Figure 7.**
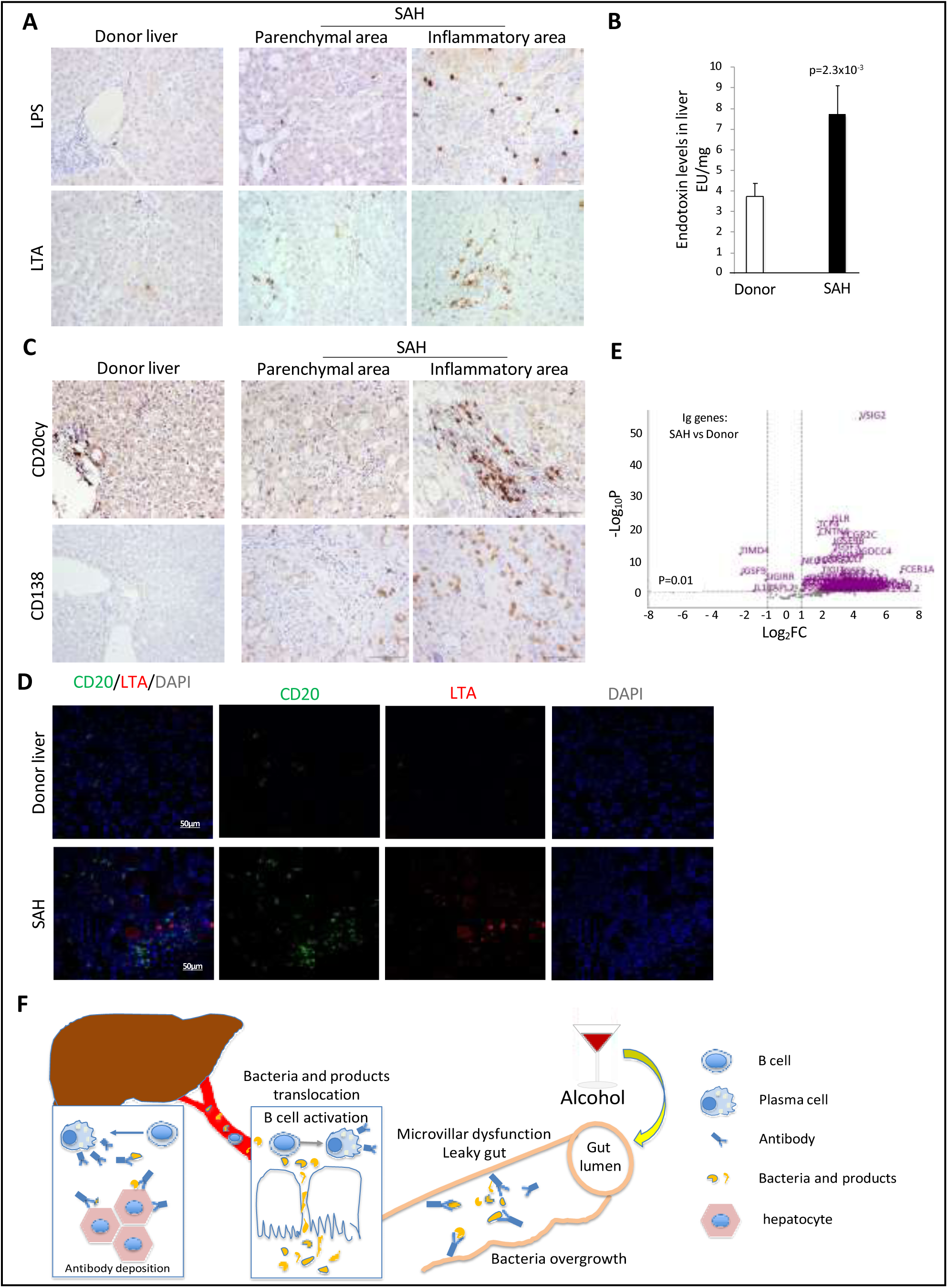
Increased bacterial antigens, B and plasma cell infiltration and immunoglobulin gene expression in SAH livers. (**A**) Immunohistochemistry staining for the gram-negative bacterial product lipopolysaccharides (LPS) and gram-positive bacteria antigen lipoteichoic acid (LTA) in SAH livers. (**B**) LPS levels in SAH liver tissues were quantified by using Pierce™ Chromogenic Endotoxin Quant Kit (n=7/group). (**C**) Immunohistochemistry staining for CD20+ B cells and CD138+ plasma cells in SAH livers. Representative images from 20 SAH or 10 HD livers (**A-C**). (**D**) Double immunofluorescence staining with CD20 (green) and LTA (red) showed colocalization of B cells and bacteria antigens in the inflammatory areas. Representative tissue sections from 6 samples per group. (**E**) RNA-seq analysis identified differentially expressed genes representing immunoglobulin fragments in SAH livers. Purple dots represented 91 differentially expressed immunoglobulin genes (log2FC >1 & adjusted p-value < 0.01) in the comparison of SAH patients between normal donors. Among them, 87 were up-regulated and 4 down-regulated. Gray dots were the remaining immunoglobulin genes that did not meet the thresholds.

Immunohistochemistry staining for CD20+ and CD138+ cells revealed that most of these cells were localized in the inflammatory areas of SAH livers, with none in HD livers (***Figure 7C***). These results suggest that increased gut bacterial antigens in the SAH liver are associated with increased B cells and plasma cells.

Based on our findings that immunoglobulins extracted from SAH livers but not present in the serum exhibited hepatocyte killing efficacy (***Figure 2G***), we suspected that immunoglobulin antibodies deposited in hepatocytes might not be produced systemically but by plasma cells infiltrating the liver. To determine if infiltration of B and plasma cells in the SAH liver is associated with expression of genes coding for certain immunoglobulins or fragments of immunoglobulins, we analyzed RNA seq data generated from SAH and healthy donor livers (GEO GSE143318). Specifically, we examined whether genes coding for immunoglobulins or fragments of specific immunoglobulins are upregulated in SAH liver samples relative to donor liver samples. We identified 175 genes coding for immunoglobulins or fragments of immunoglobulins and compared their expression levels between SAH and control samples.

Interestingly, 87 identified genes representing immunoglobulin fragments showed a robust and stable expression in SAH livers but not in healthy donor livers (***Figure 7D***). These results suggest that infiltrating B and plasma cells secrete immunoglobulins that not only have anti-intestinal antigen activity but also cross-react with hepatocyte antigens.

## Discussion

Using explanted livers from SAH patients, we discovered massive antibody deposition in ballooned hepatocytes which is associated with complement activation. Antibodies from the SAH liver but not patient serum exhibited hepatocyte killing efficacy in vitro. Unique antibodies found in the SAH liver not only recognize bacterial (*E. coli*) antigens but also cross-react with a large number of human antigens. Our data support the hypothesis that anti-bacterial antibodies and/or plasma cells originating from gut translocate to the liver due to gut leakiness caused by excessive alcohol drinking and that these hepatic plasma cells produce antibodies which not only recognize intestinal bacterial antigens but also cross react with antigens of human hepatocytes (***Figure 6E***). Antibody deposition, with complement activation as well as immune cell activation, may result in acute liver failure due to antibody-mediated inflammation.

Leaky gut, also known as increased intestinal permeability, is a digestive condition in which bacteria and toxins are able to pass from the gastrointestinal tract to extraintestinal sites and facilitate an increased ingress of inflammatory cytokines and endotoxin to the liver^22^. Unique anti-*E. coli* IgG or IgA antibodies are identified in alcoholic livers, but not other liver diseases (***table supplement 2***) suggesting the impact of excessive alcohol drinking on translocation of anti-bacterial antibodies to the liver. Interestingly, few anti-bacterial antibodies in AC livers cross-reacted with human antigens (***figure supplement 4***), while a large number of anti- bacterial antibodies in SAH livers also recognized human antigens. The differing gut microbiota between actively drinking patients with cirrhosis and those with alcoholic hepatitis^23,24,25^ supports the notion that alcohol-induced changes of gut microbiota composition may contribute to the development of anti-bacterial antibodies that cross-reacted with human antigens. SAH is an acute-on-chronic liver failure. Cross-reacting anti-bacterial autoantibodies translocated to the liver may determine the progress from chronic liver disease toward acute hepatitis.

Coincidentally, we found a significant number of unique IgA autoantibodies in HBV and unique IgG autoantibodies in HCV livers (***table supplement 1***) suggesting these IgA or IgG autoantibodies deposited in the liver might be derived from systemic autoimmune process which is ongoing in HBV or HCV infected patients^26, 27^. Previous studies have clearly established the co-occurrence of certain autoantibodies in patients with chronic HBV or HCV infection and the autoantibody positivity was common in HBV and HCV patients. A strong association between increasing serum IgA level and disease progressing in patients with chronic HBV infection has been reported^26^. Similarly, serum IgG was increased in patients with chronic hepatitis C with respect to both alcoholics and healthy controls^27^. No unique anti-*E. coli* protein IgA or IgG antibody was identified in HBV or HCV livers, indicating autoantibody production was related to the virus rather than bacterial infection.

The presence of cross-reacting IgM antibodies against *E. coli* and human antigens in the PBC livers is worth noting. Current data suggest that PBC is an autoimmune disease^28, 29^ and an infectious etiology as trigger for development of PBC has been postulated. Antibodies reacting against the mitochondrial human pyruvate dehydrogenase complex which are the serologic hallmark of PBC cross react with the *E. coli* pyruvate dehydrogenase complex, implicating *E. coli* infection in the pathogenesis of PBC^30^. Interestingly, patients with PBC had a much higher number of inducible IgM-producing B-cells in peripheral blood and each B-cell produced a greater quantity of IgM protein compared with control^29^. A large number of unique human and *E. coli* antigens were recognized specifically by IgM from the PBC livers, supporting the theory of bacterial infection-related IgM production in pathogenesis of PBC.

Our findings suggest that SAH may be an antibody-mediated disease due to intrahepatic etiology. Unlike antibody mediated autoimmune disease where systemic B cells produce autoantibodies against self-antigens and unlike the fact that autoimmune diseases can be transferred from an affected patient to a normal individual by the transfer of patient-derived serum (or immunoglobulins), serum from SAH patients did not show hepatocyte killing efficacy in vitro. Importantly, no antibody-mediated rejection was observed in SAH patients following liver transplantation^14^. It is perhaps more likely that gut-derived plasma cells were resident in SAH livers, and these plasma cells produce anti-bacteria antibodies which strongly cross-react with hepatocyte antigens. The diseased liver absorbs these gut-derived and locally produced antibodies without releasing them to the circulation. With the functional deposition of antibodies in the SAH liver (***Figure 2***), activation of complement through the classical pathway which has been reported previously^31^ leads to ballooning degeneration of hepatocytes which is the predominant mode of injury in alcoholic hepatitis and untreatable inflammation. When antibody is deposited, it may also impair a number of important biological process in hepatocytes (***Figure 6***). This pathophysiology implies severe damage from short-lived leaky gut due to a transient interval of high alcohol intake with ongoing residual continued damage due to the antibody production that continues only within the liver to be replaced. Therefore, it is reassuring that the new transplanted liver will not be damaged because the pathogenesis, involving the consequences of leaky gut pathology, will end quickly as long as the recipient remains abstinent, by which the new liver is not threatened by the SAH process.

The main limitations of the present study are the limited sample size, the lack of whole gut bacterial-proteome microarrays and the lack of animal models of severe alcoholic hepatitis. Nevertheless, identifying the presence of anti-bacterial autoantibodies in SAH livers provides a new insight into pathogenesis of SAH. Future studies utilizing needle biopsy samples from mild and moderate alcoholic hepatitis patients could assess infiltration of B and plasma cells and a likely correlation between antibody deposition and severity of hepatitis. For example, infiltration of plasma cells and antibody deposition may predict the progression toward SAH.

This may be a new therapeutic strategy in alcoholic hepatitis patients. First, preventing gut leakiness caused by alcohol drinking may prevent SAH. Second, eliminating antibody producing B/plasma cells in the liver by using anti-CD20 may serve. Finally, a trial using complement inhibitors such as Eculizumab may be worth considering.

## Methods

### Collection of liver tissue samples

Explanted liver tissues and blood were collected from patients with SAH or AC who were referred for liver transplantation at Johns Hopkins hospital^14^. Explanted liver tissues from patients with other liver diseases were obtained from the Liver Tissue Procurement and Distribution System at the University of Minnesota, which was funded by NIH Contract# HHSN276201200017C. All studies were approved by the Johns Hopkins Medicine Institutional Review Boards (IRB00107893).

### Immunohistochemistry

Cut sections were prepared from formalin-fixed paraffin embedded liver tissues for staining with IgG (ab200699), IgA (ab200699 or GTX20770), C4d (ab167093), C3d (ab136916), CD20 (Dako, Santa Clara, CA), CD138 (Abcam, Cambridge, MA), *E. coli* LPS (Abcam, Cambridge, MA), LTA (Thermo Fisher, Waltham, MA). Vectastain Elite ABC Staining Kit and DAB Peroxidase Substrate Kit (Vector Laboratories, Burlingame, CA) were used to visualize the staining according to the manufacturer’s instructions. Diaminobenzidine tetrahydrocholoride and blue alkaline phosphatase (Vector Laboratories) were used as brown and blue chromogen and hematoxylin as nuclear counterstaining. Echo Revolve microscope (Echo Laboratories Inc) was used for taking image pictures.

### ELISA

Liver protein lysates used for this assay contained similar concentrations of protein. Each SAH and donor liver sample were added on a precoated ELISA plate (Thermofisher Scientific, Waltham, MA) to determine the total IgG (BMS2091), IgA (BMS2096), IgM (BMS2098) and IgG subclasses (991000).

### Isolation of Peripheral Blood Mononuclear Cells (PBMCs)

PBMCs were isolated from heparinized peripheral blood samples from healthy volunteers using Ficoll-Paque plus density gradient medium.

### Antibody-dependent cell-mediated cytotoxicity (ADCC) assay

ADCC was determined by a calcein-acetyoxymethyl release assay (calcein-AM, C3100MP, Thermofisher Scientific). Calcein-AM labeled primary human hepatocytes were cultured with Ig from SAH livers, patient serum or human IgG isotype control in a 96 well plate at a density of 1x10^4^ cells per well in triplicate and PBMCs were added as effector cells at an effector: target cell ratio of 5:1 respectively. Antibody-independent cell mediated cytotoxicity (AICC) was measured in wells containing target and effector cells without the addition of AH or control IgG antibodies. The following formula was used to calculate ADCC: % ADCC= 100 x (mean experimental release –mean AICC) ÷ (mean maximum release – mean spontaneous release).

### Western blot analysis

Fifty-milligram liver tissues from SAH or the controls were homogenized in the lysis buffer (Cell Signaling Technology, MA). The total protein concentrations of each liver samples were determine using a standard curve generated with BSA at different known concentration using the Quick Start™ Bradford Protein Assay (Bio-Rad, U.S.A.). On the basis of the measure protein cencetrions, 25 micrograms of total proteins of each liver sample were boiled in NuPAGE LDS Sample Buffer (Thermofisher, MA) and subjected to electrophoresis in a 4-12 % Bis-Tris gradient PEG gel (Thermofisher, MA). All samples were tested in several parallel gels. One gel was stained with SimplyBlue™ SafeStain according to the product manual (Thermofisher, MA), and the other one was subjected to the western blot assay using the Trans-Blot Turbo RTA Midi 0.45 µm LF PVDF Transfer Kit (Bio-Rad Laboratories, CA). After transferring the total proteins to the PVDF membrane, the membranes were incubated with the IRDye labeled human immunoglobulin antibodies specific recognizing IgG, IgM, IgA, and IgE, or mouse monoclonal antibodies against human C3d (Bio-Rad Laboratories, CA) and C4d (Santa Cruz, CA) respectively, followed by probing with Alexa 647 labeled Goat anti-Mouse IgG (H+L) (Thermofisher, MA). After scanning with Odyssey CLx Imaging System, the signals were calculated by the corresponding software and then analyzed by Excel.

### Multiplex immunofluorescence staining

Sequential multiplex immunofluorescence staining on formalin-fixed, paraffin-embedded liver sections was performed, as previously described^32^. Images were acquired on a Zeiss LSM 900 confocal microscope. Acquired images were processed and analyzed using FIJI^33^. The following antibodies were used: HepPar1 (Catalog NBP2-45272, Novus), CD68 (Catalog M0876, Dako), CD32 (Catalog 53151, Cell Signaling Technology), IgG (109-005-088, Jackson ImmunoResearch), IgA (Catalog ab124716, Abcam), β-catenin (Catalog 610154, BD Biosceinces), C4d (Catalog BI- RC4D, BIOMEDICA).

### Antibody extraction and protein microarray analysis

The liver tissues were subjected to a Dounce homogenizer in a lysis buffer (0.1 M Glycine pH 2.0, 150 mM NaCl) to elute the binding antibodies (Ig). After a five-minute spin down with 20,000 g at 4°C, the supernatants were transferred to new 15ml tubes and neutralized with 1M Tris-base buffer to pH 7.0 immediately. Then, the antibody enrichment was performed using Protein L coupled magnetic beads (Thermofisher, MA) according to the manual. The antibodies extracted from 4.8-gram liver tissue pieces were subjected to the protein microarray assays using human proteome microarray HuProt^®^ array and *E. coli* strain K-12 bacterial proteome microarray respectively to screen their corresponding antigens^15, 16^. Data analysis including the criteria for positive hits was performed as before^15, 16^.

### Identification of cross-reactive antibodies against human and bacterial proteins

The total lysates of *E. coli* were immobilized on the magnetic beads using the kit of Pierce NHS-Activated Magnetic Beads according to the product manual (ThermoFisher, MA). Then, these E. coli protein magnetic beads were incubated with the extracted total autoantibodies from liver samples to capture the active antibodies. After eluting with Glycine pH 2.0, the eluted antibodies were neutralized with 1 M tris-base buffer to pH 7.0 immediately. Finally, these antibodies were subjected to the human proteome microarray.

### RNA-sequencing and data processing

The total RNA was isolated using Qiagen RNeasy© kit. After the RNA quality was assessed by capillary electrophoresis (bioanalyzer), cDNA libraries were prepared using TruSeq© RNA Library Prep Kit and sequenced with an Illumina NextSeq500. Base-calling and fastq conversion was performed using RTA (2.4.11) and Bcl2fastq (2.18.0.12), respectively. Raw sequencing files were uploaded to the NIH GEO database.

Adaptor sequences were trimmed from the raw reads using Cutadapt^17^. Trimmed reads were then mapped to reference genome GRCh38 using STAR aligner with default parameters^18^. The number of counts per gene was estimated using the “quantMode” command in STAR. Batch effect was corrected using Combat seq. Differentially expressed genes (DEGs) were then identified using DESeq2 ^19^. Genes with adjusted *P* < 0.01 and log2 fold change > 1 were chosen as DEGs.

### Gene Ontology (GO) analysis

DAVID^20^ was used to conducted GO analysis to find out enriched GO terms (cellular components and biological process). All enriched terms were chosen with a threshold p-value of 0.05.

### Data availability

External data requests can be directed to the corresponding authors, who will respond within 1 week and help facilitate the request. Access to clinical datasets will be available based on approval through the IRBs of the Johns Hopkins University and might be subject to patient privacy.

## Acknowledgments

The authors thank patients and liver donors, their families and surrogates and medical personnel. We thank the anesthesiologists, nurses and transplantation fellows at the Johns Hopkins hospital who assisted in collecting samples. In addition, we thank clinical pathology core at Johns Hopkins hospital for performing immunohistochemistry staining and scanning the slides.

## Additional Information

### Author contributions

A.R.A. and Z.S. conceived the project. Z.S. and H.Z. designed all experiments. A.M.C, R.W., A.G., B.P, S.O, B.K., and R.A.A. enrolled patients and collected tissue samples for the study. A.R.A, J.M., L.Q., B.P. performed immunohistochemistry staining, ELISA, ADCC assays, RNA sequencing and data analysis. G.S. and H.Z. performed western blot analysis, HuProt and E. coli proteome arrays. C.S.C. provided protein chips for E. coli arrays. T.G., X.H., M.H. and J.Q. designed data and statistical analysis for proteome arrays and RNA sequencing. Z.Z. performed LPS assays. D.F. and H.J.L performed multiplex immunofluorescence staining. A.M.C., B.G., J.B. and J.K.B. provided intellectual input. Z.S. wrote and A.R.A., H.Z., J.Q., B.G., A.M.C and J.K.B. edited the manuscript. All authors reviewed the manuscript.

### Ethics

All studies were approved by the Johns Hopkins Medicine Institutional Review Boards (IRB00107893 and IRB00154881).

## Additional files

### Figure supplements 1-6

**Figure S1.**
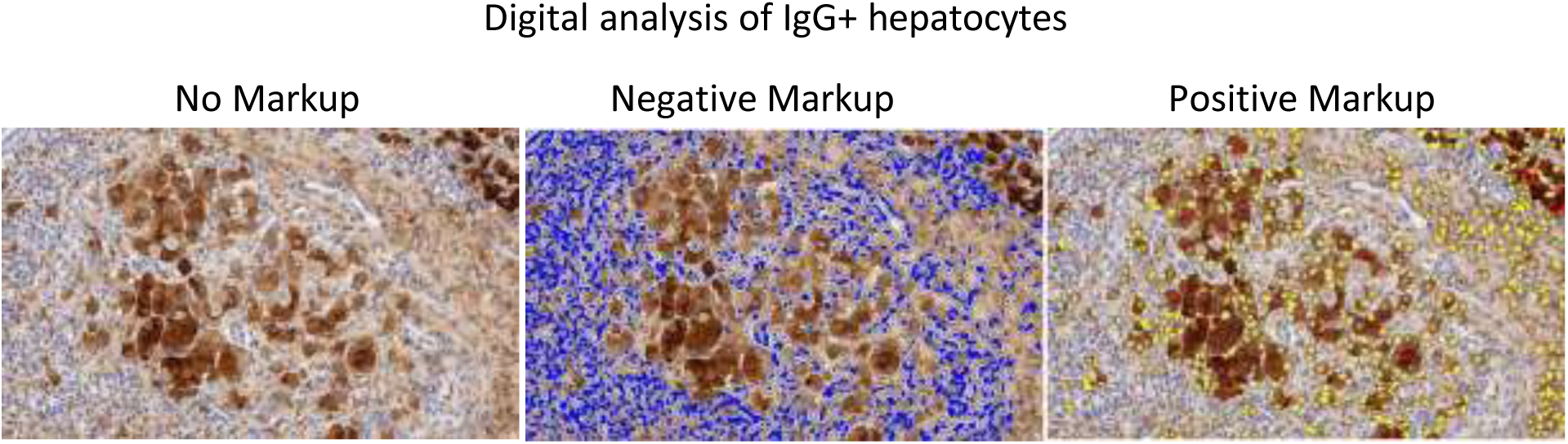
HALO image analysis of IgG positive hepatocytes in severe alcoholic hepatitis livers. Representative images of liver tissue sections stained with anti-human IgG antibody. Positive cells were reported as percentage stained surface area of total annotated area by digital analysis (Hallo, Indicalabs, Corrales, NM).

**Figure S2.**
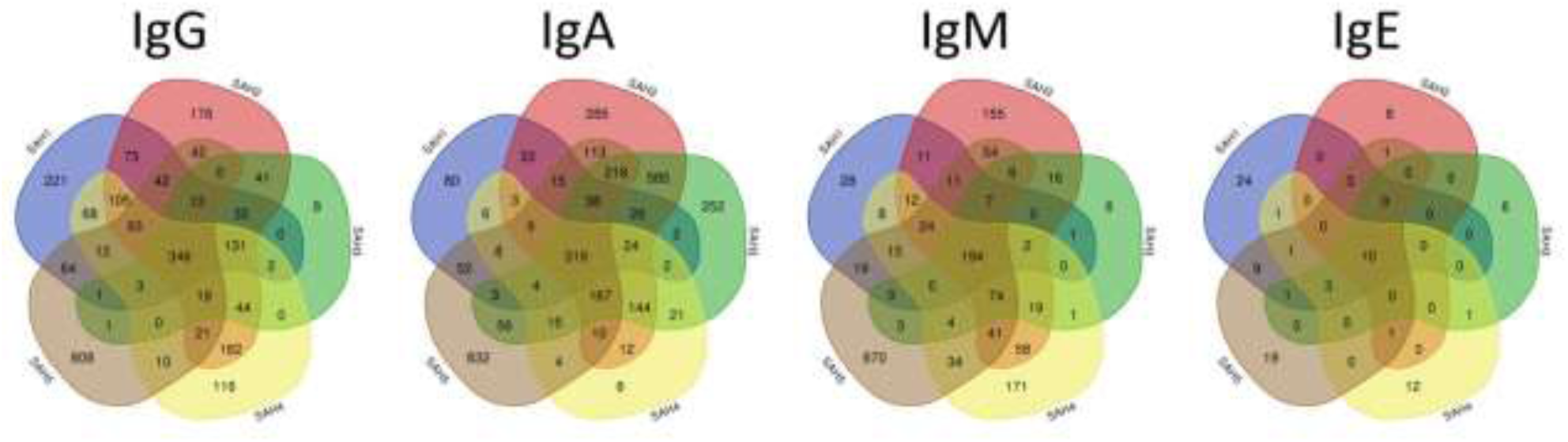
Venn diagram analysis of autoantigens recognized by antibodies extracted from the five SAH liver tissues. After individual elution of antibodies from the five SAH liver tissues (i.e., SAH1-5), they were individually profiled on the HuProt arrays. The identified autoantigens were grouped on the basis of anti-IgG, -IgA, -IgM, and -IgE isotypes, and the shared and unique autoantigens were analyzed using the Venn diagram analysis. As illustrated in each isotype panel, the shared autoantigens recognized by the anti-IgG, -IgA, -IgM, and -IgE antibodies among the five SAH samples are 346, 319, 194, and 10, respectively.

**Figure S3.**
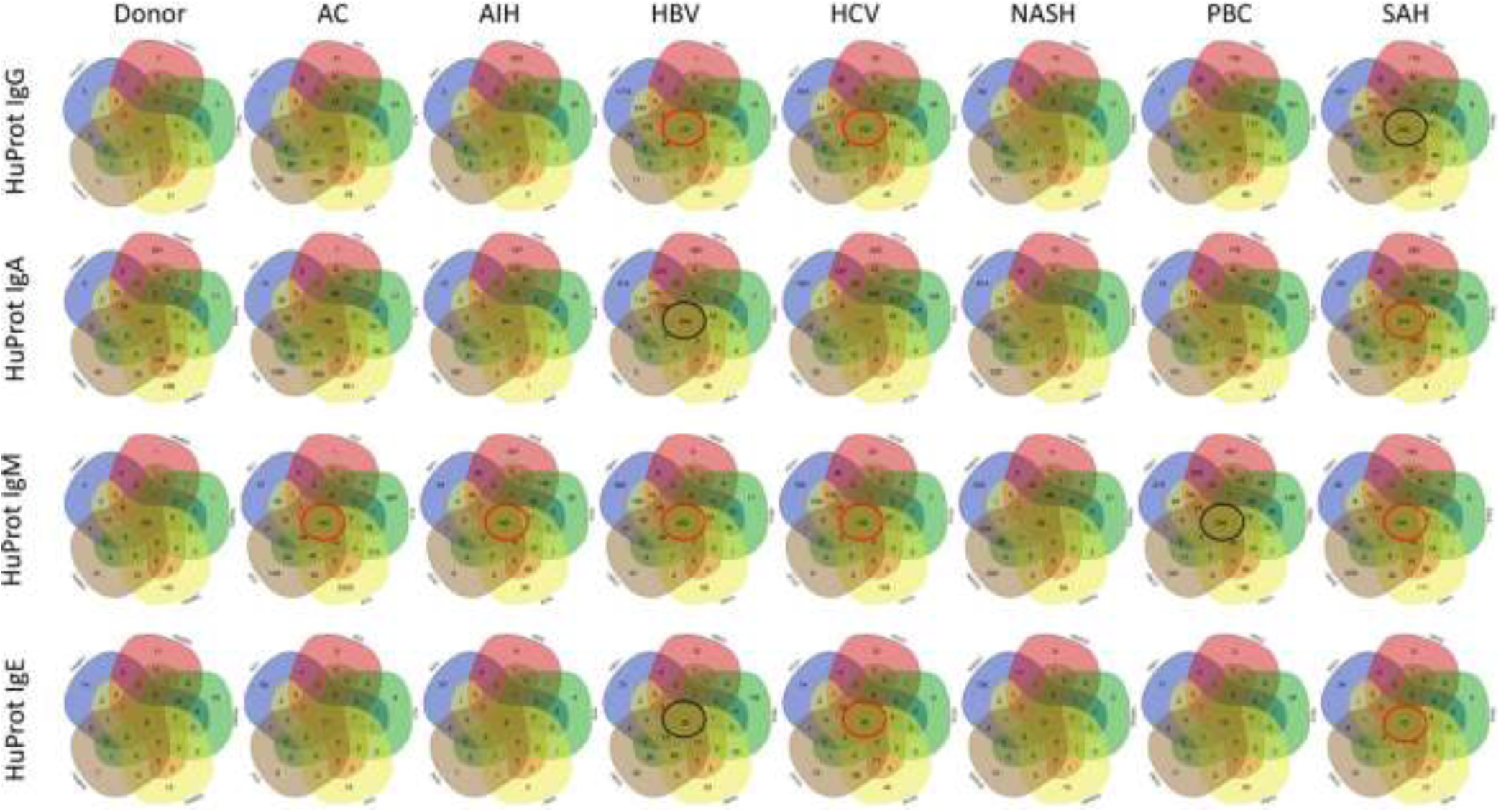
Summary of the Venn diagram analysis of autoantigens recognized by antibodies extracted from the five donor, five AC, five AIH, five HBV, five HCV, five NASH, five PBC, and five SAH liver tissues. Using the same approach as described in Figure S2, the shared and disease-specific autoantibodies were identified. The Ig isotypes are designated in each roll of the Venn diagrams, and the tissue types are shown on the top of each column of the Venn diagrams.

**Figure S4.**
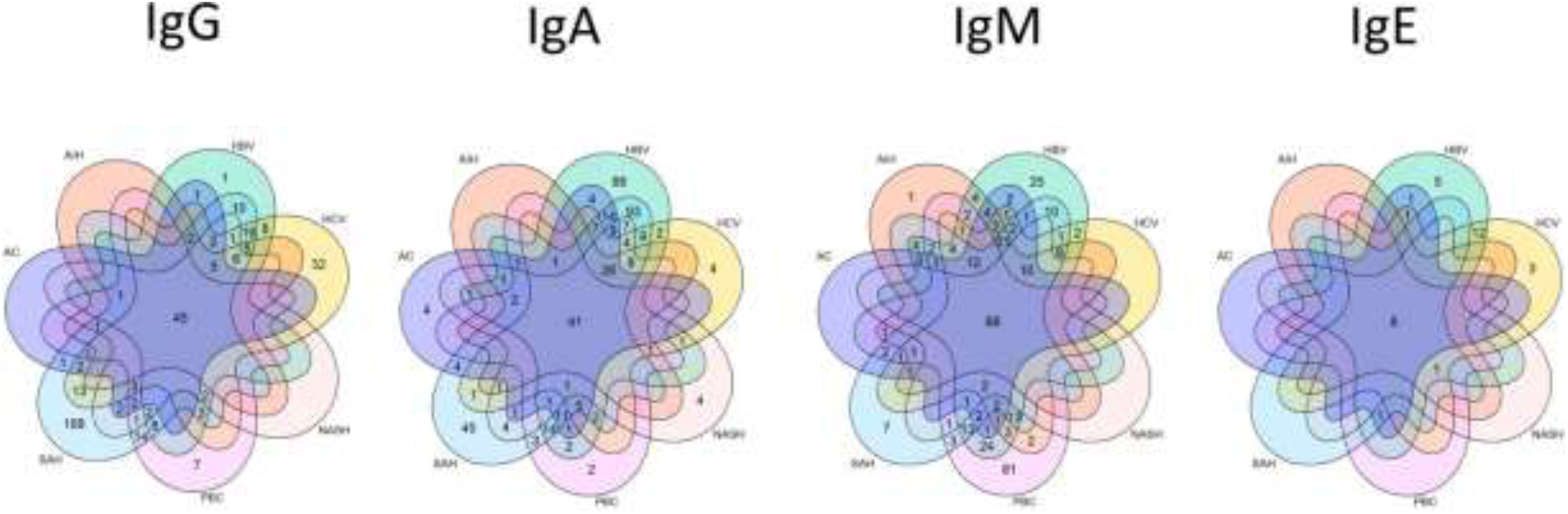
A seven-member Venn diagram analysis of the shared autoantigens recognized by antibodies extracted from the diseased liver tissues. The shared autoantigens identified by the antibodies extracted from the seven diseased liver tissues were subjected to a seven-member Venn diagram analysis. The Ig isotypes are shown on the top of each Venn diagram.

**Figure S5.**
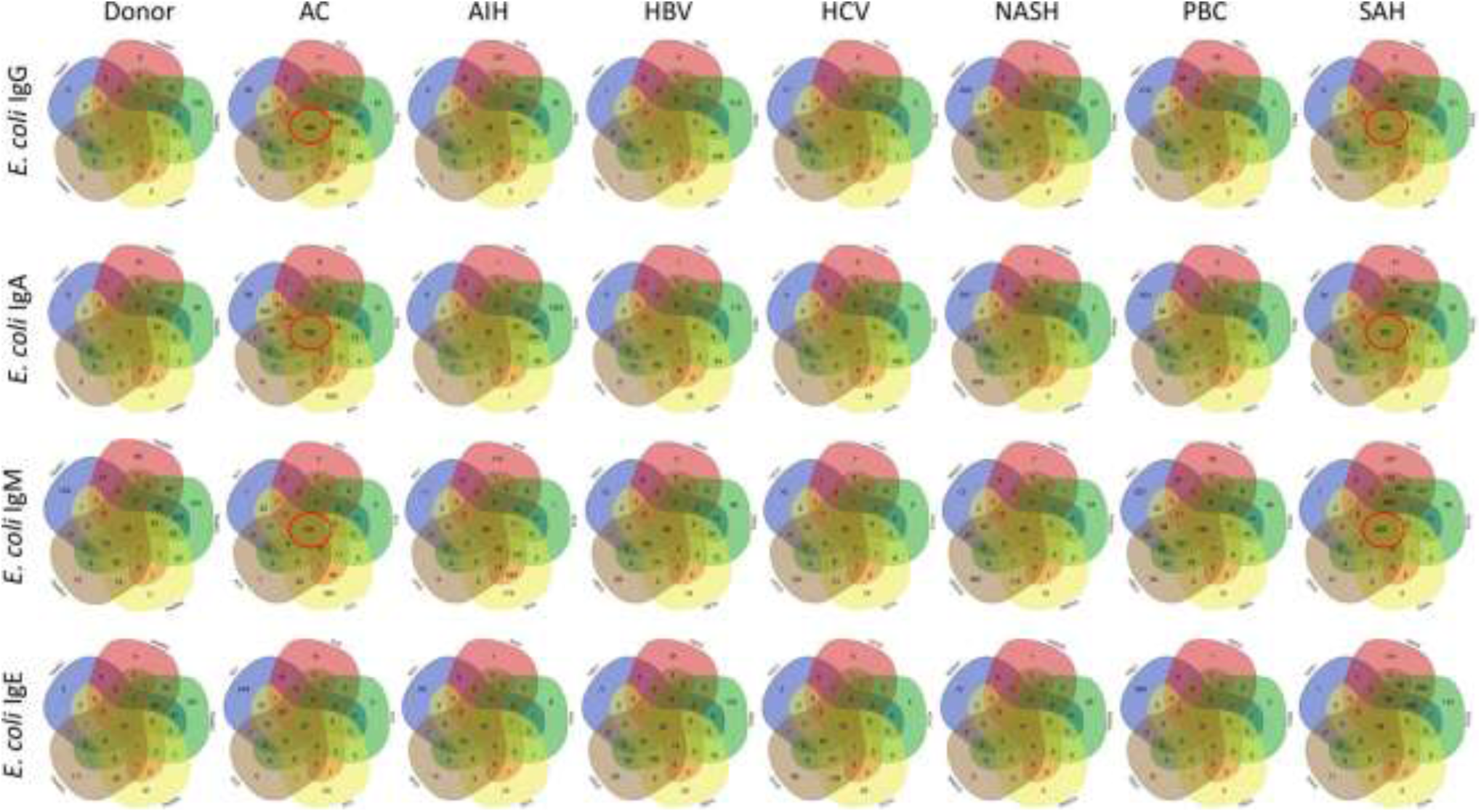
Summary of the Venn diagram analysis of bacterial antigens recognized by human antibodies extracted from the five donor, five AC, five AIH, five HBV, five HCV, five NASH, five PBC, and five SAH liver tissues. Using the same approach as described in Figure S1, the shared and disease-specific antibodies were identified. The Ig isotypes are designated in each roll of the Venn diagrams, and the tissue types are shown on the top of each column of the Venn diagrams.

**Figure S6.**
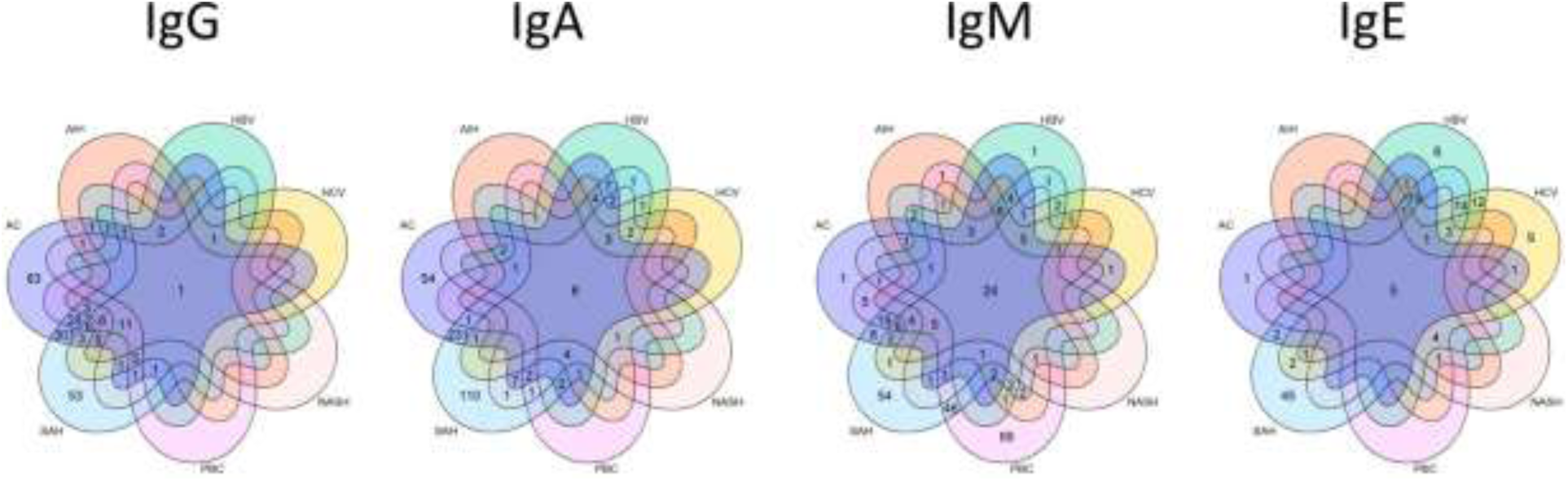
Venn diagram analysis of the shared bacterial antigens recognized by human antibodies extracted from the diseased liver tissues. The shared bacterial antigens identified by the antibodies extracted from the seven diseased liver tissues were subjected to a seven-member Venn diagram analysis. The Ig isotypes are shown on the top of each Venn diagram.

### Table supplements 1-5

**Table S1.** The numbers of unique autoantigens recognized by antibodies extracted from the diseased liver tissues (HuProt arrays).

|  | IgG-binding | IgA-binding | IgM-binding | IgE-binding | Total |
| --- | --- | --- | --- | --- | --- |
| SAH | 188 | 45 | 7 | 0 | 240 |
| AC | 0 | 4 | 0 | 0 | 4 |
| HBV | 1 | 88 | 25 | 5 | 119 |
| HCV | 32 | 4 | 0 | 3 | 39 |
| PBC | 7 | 2 | 81 | 0 | 90 |
| NASH | 0 | 4 | 0 | 0 | 4 |
| AIH | 0 | 0 | 1 | 0 | 1 |

**Table S2.** The numbers of unique *E. coli* antigens recognized by antibodies extracted from the diseased liver tissues (*E. coli* proteome arrays).

|  | IgG-binding | IgA-binding | IgM-binding | IgE-binding | Total |
| --- | --- | --- | --- | --- | --- |
| SAH | 53 | 110 | 54 | 45 | 262 |
| AC | 83 | 54 | 1 | 1 | 139 |
| HBV |  |  | 1 | 6 | 7 |
| HCV |  |  |  | 9 | 9 |
| PBC |  |  | 69 |  | 69 |
| NASH |  |  |  |  |  |
| AIH |  |  |  |  |  |

**Table S3.**
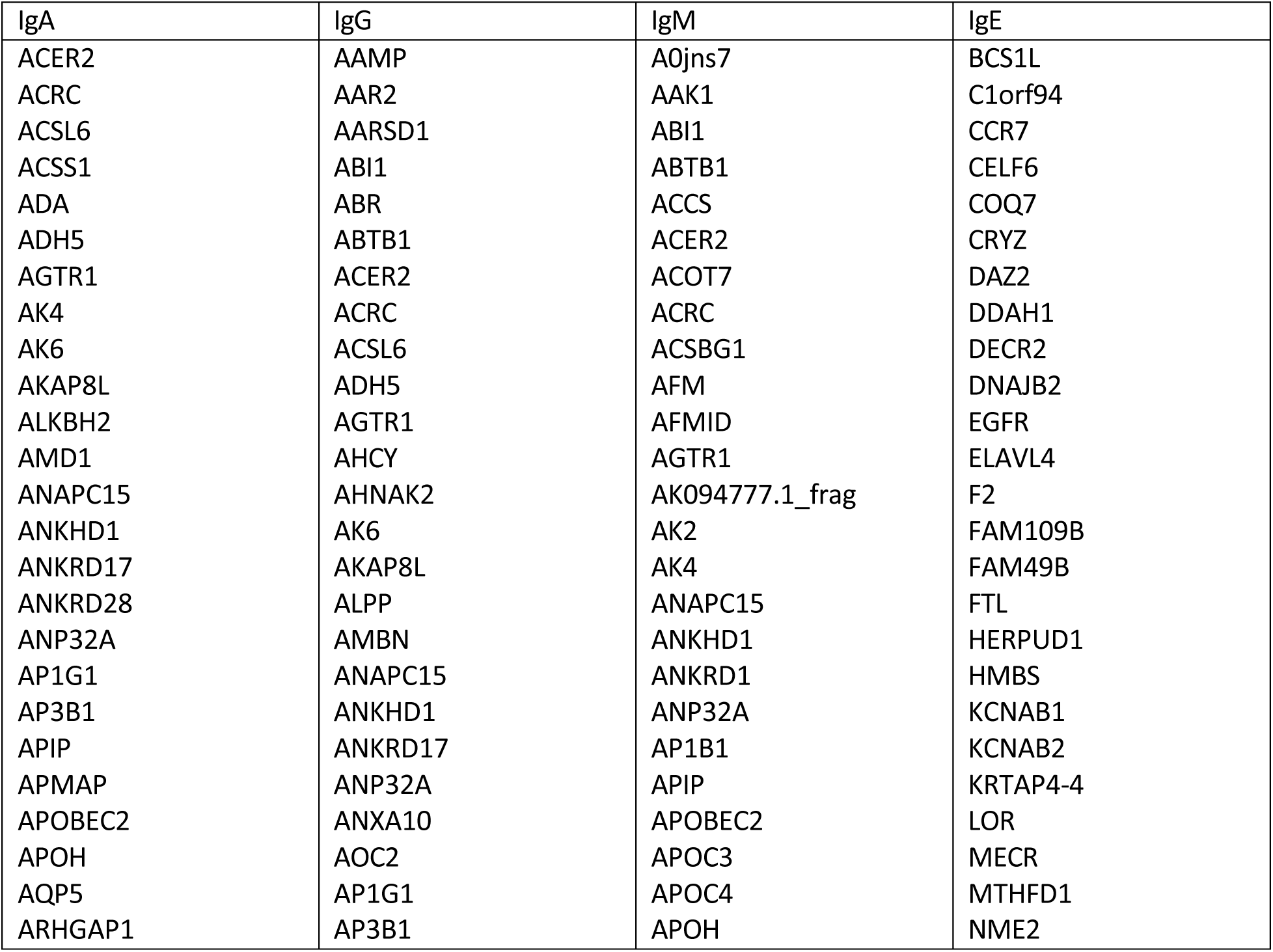
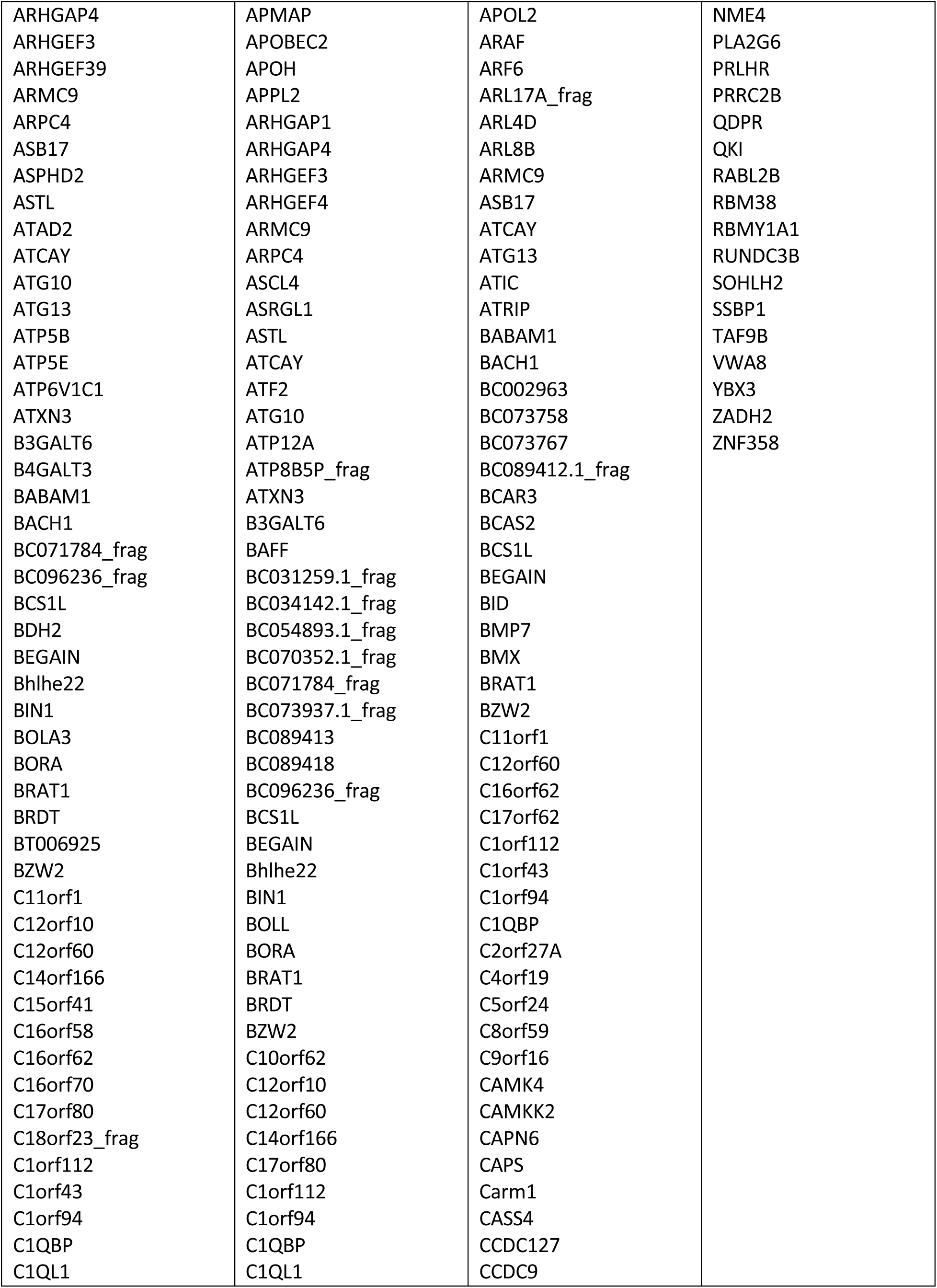
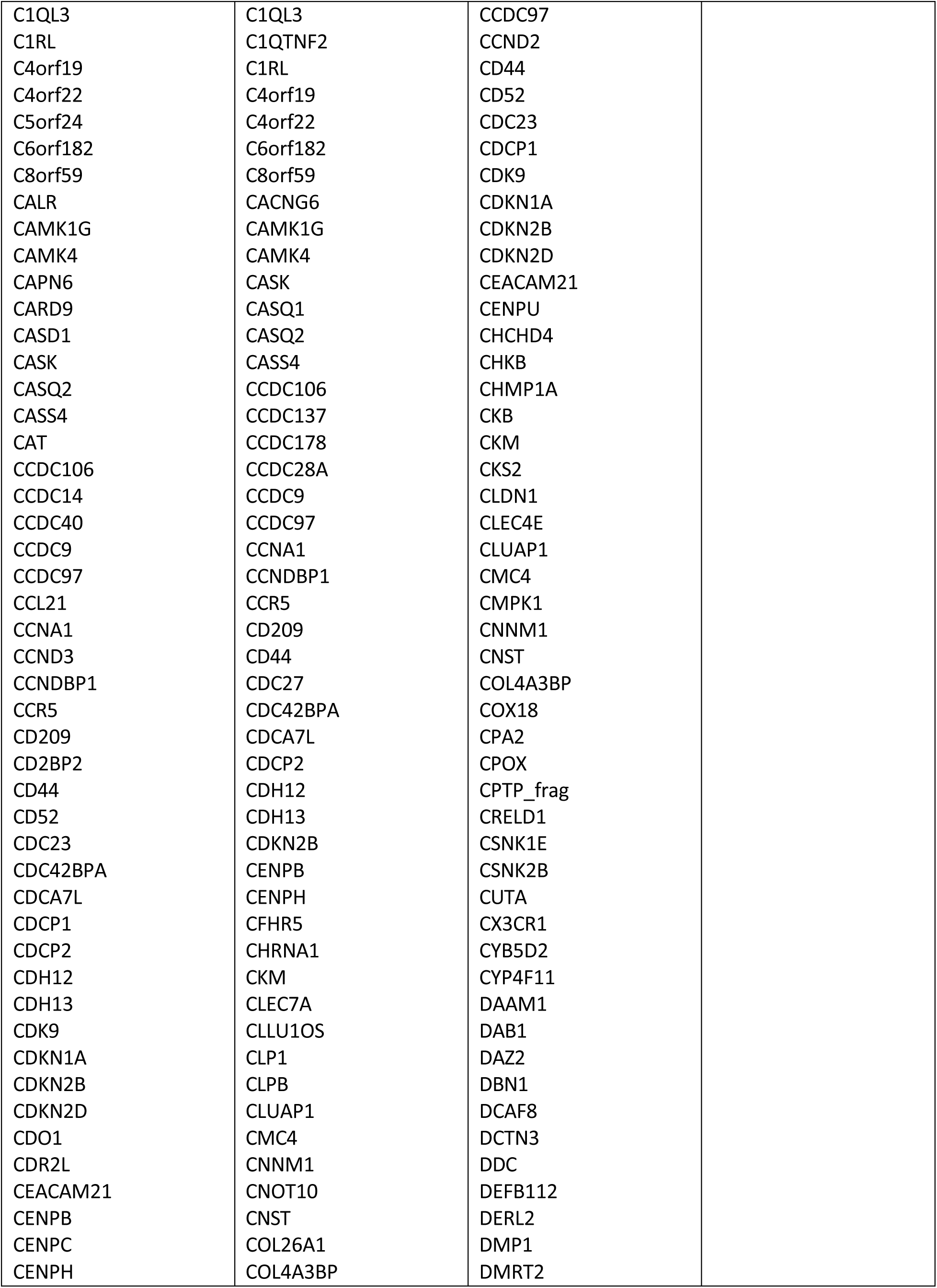
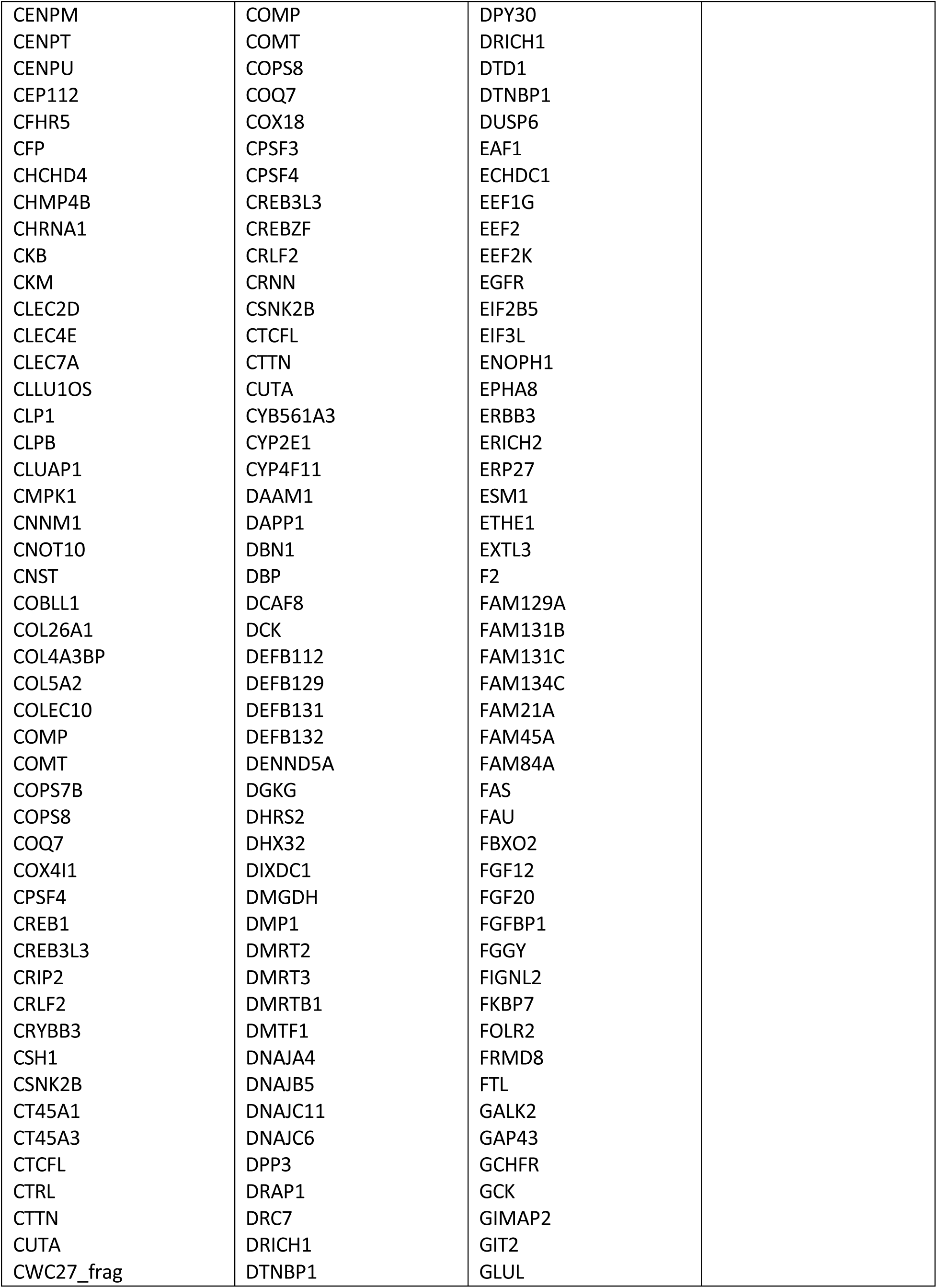
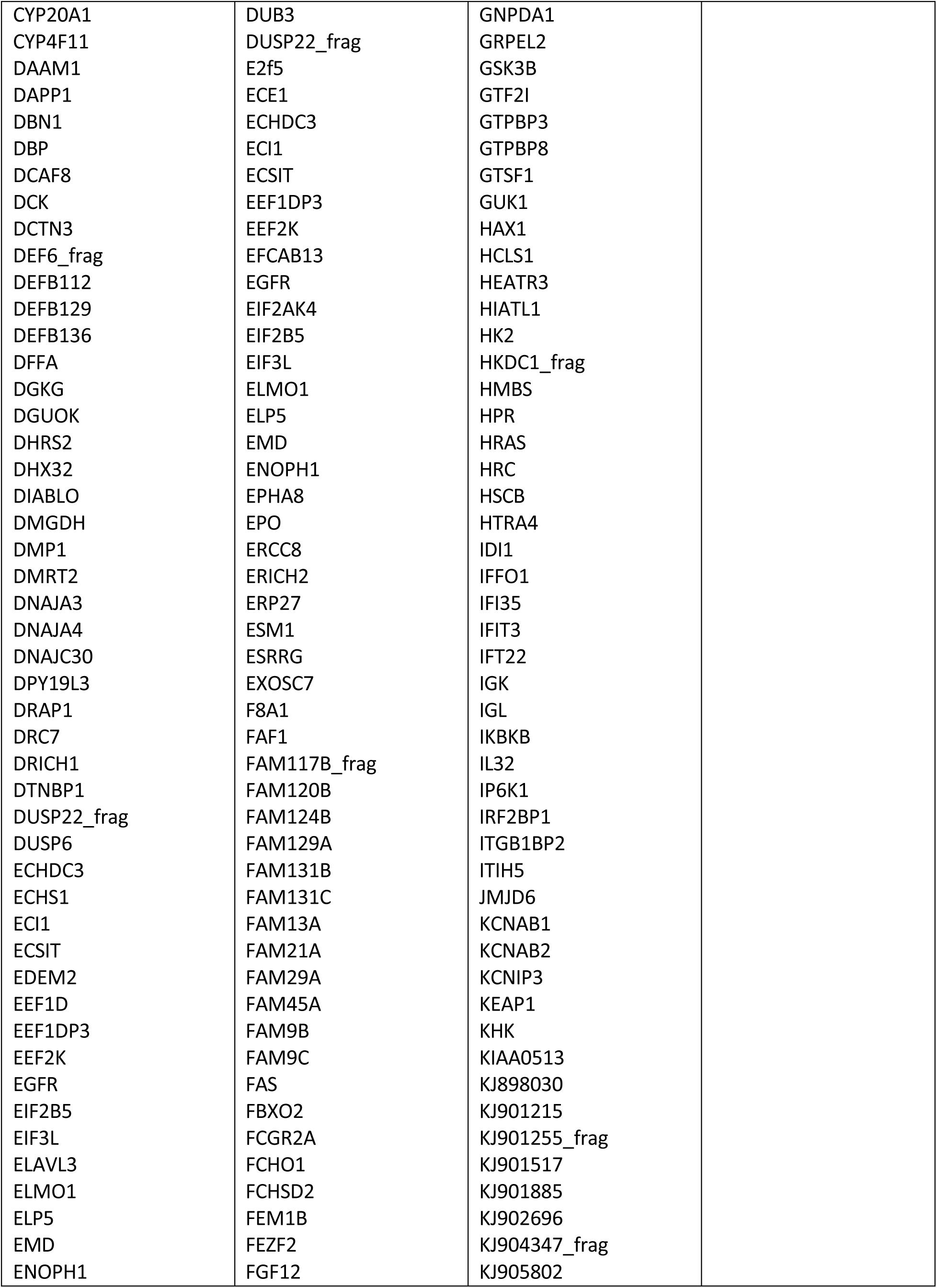
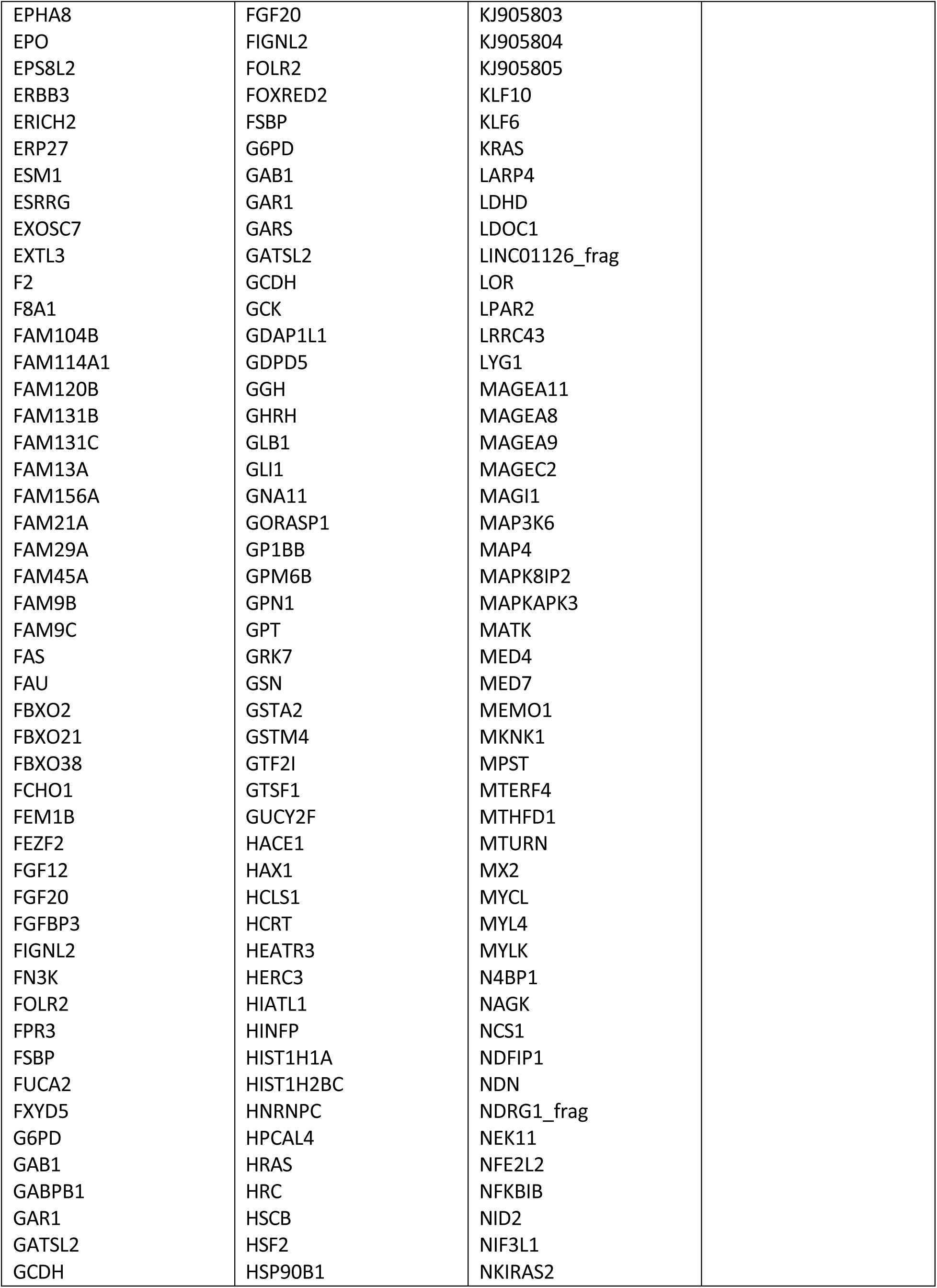
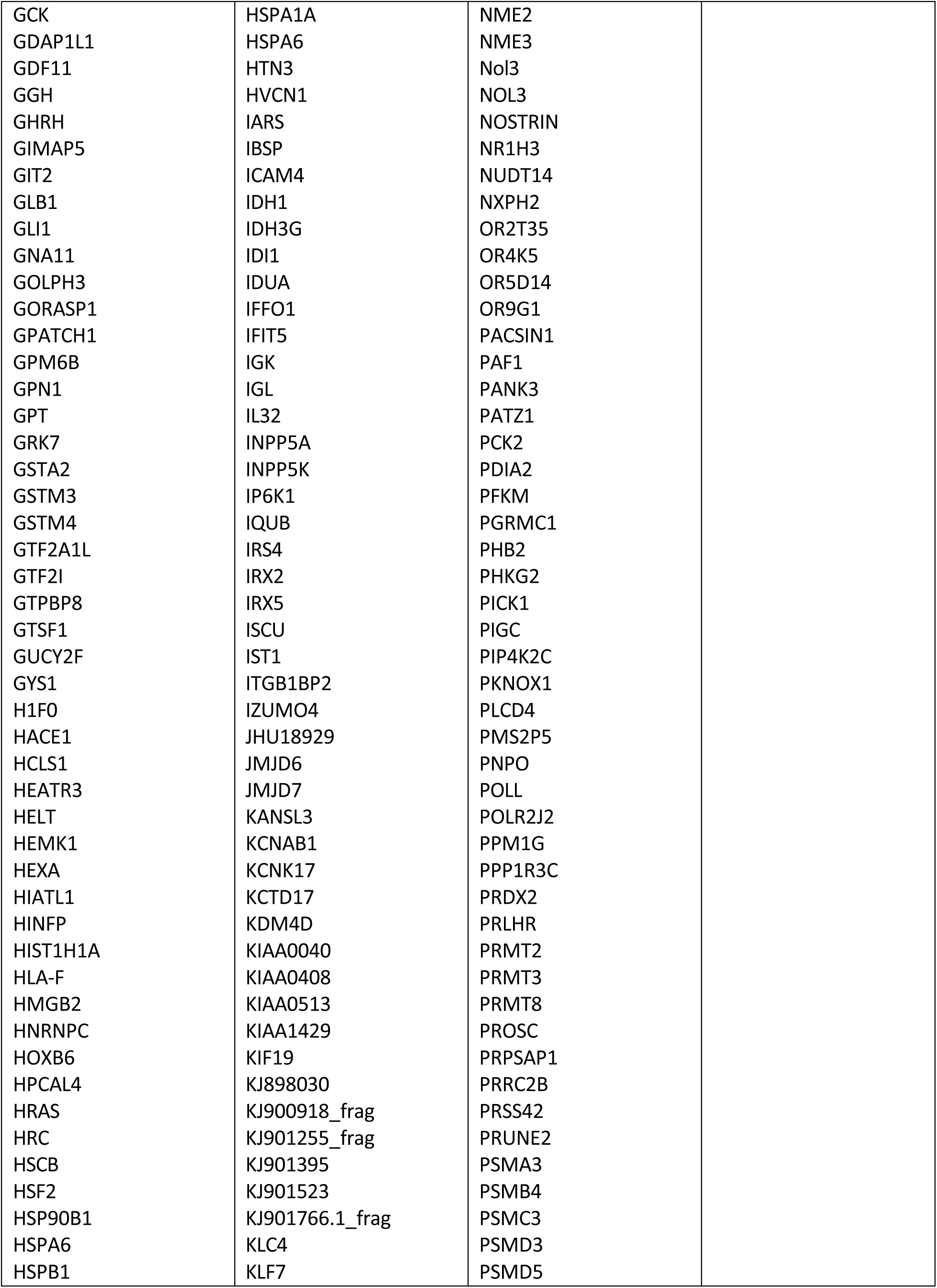
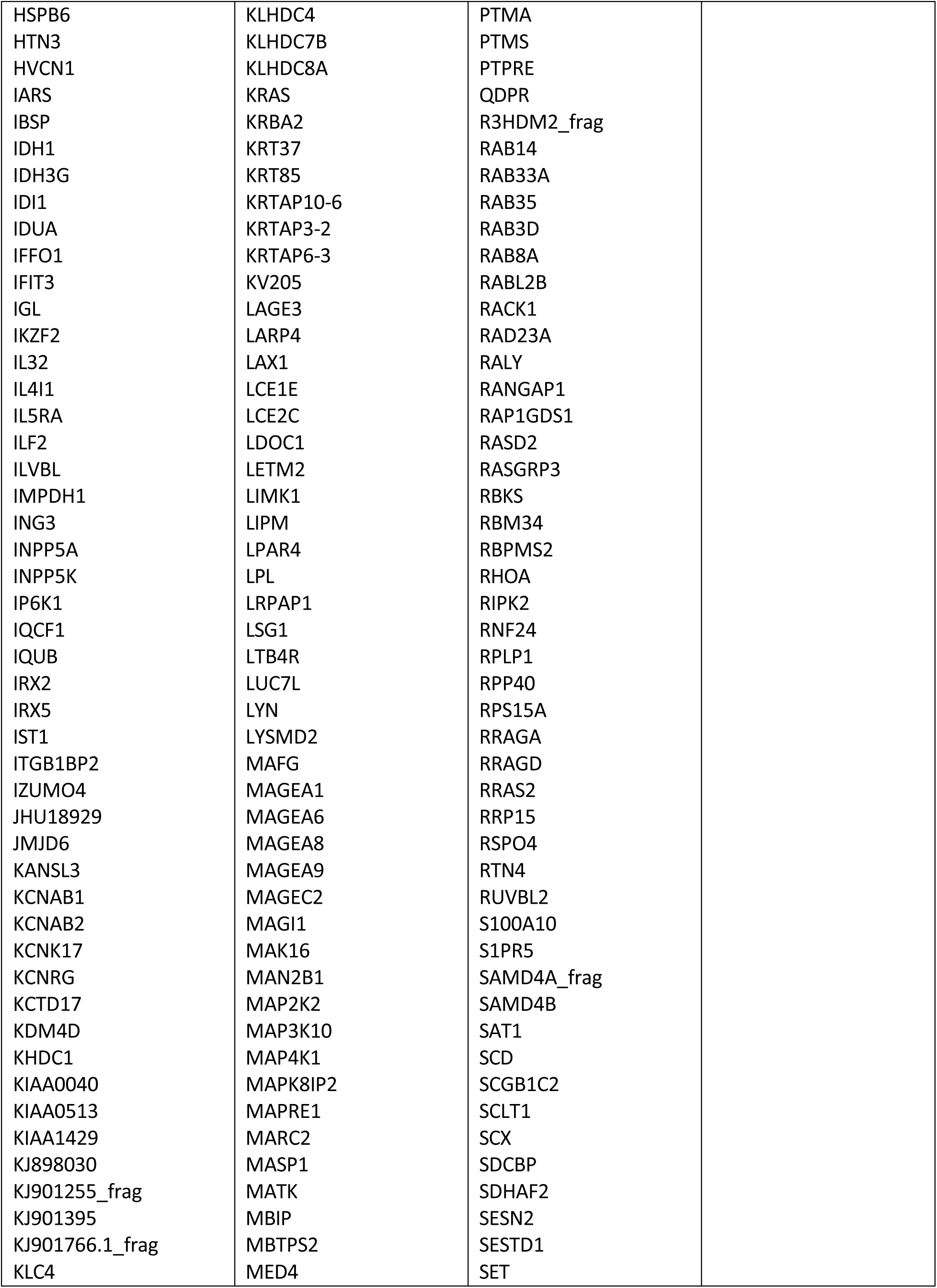
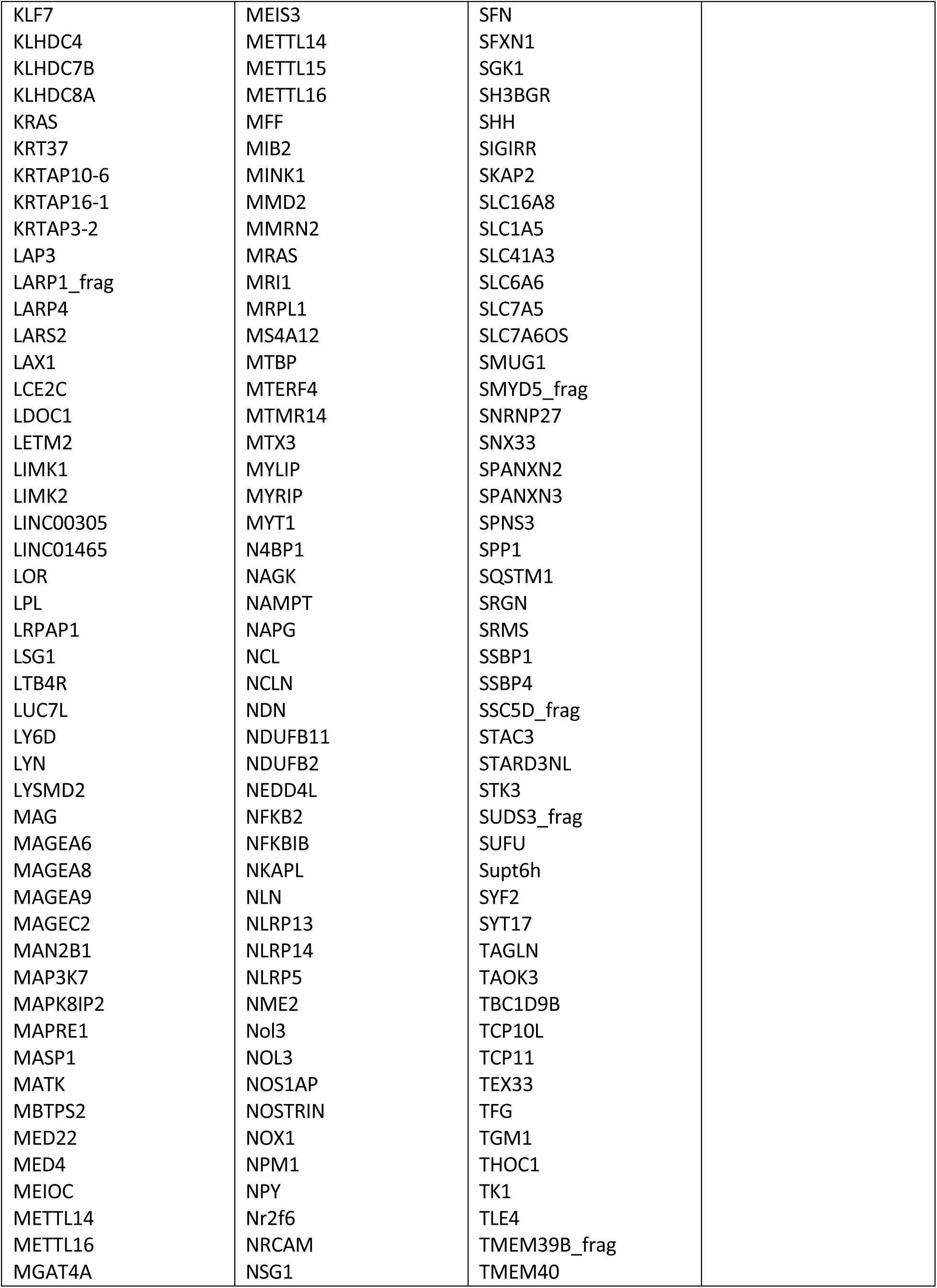
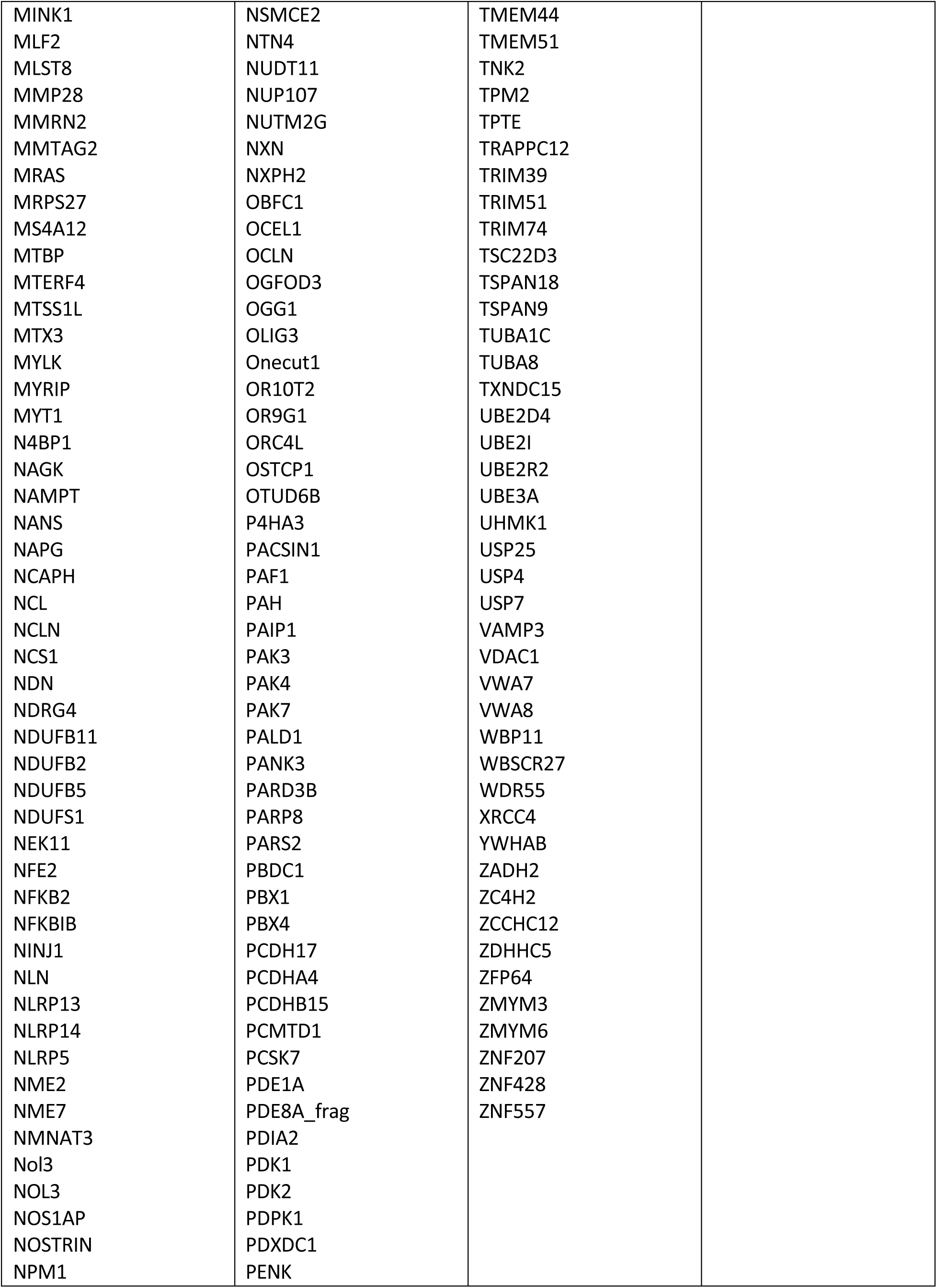
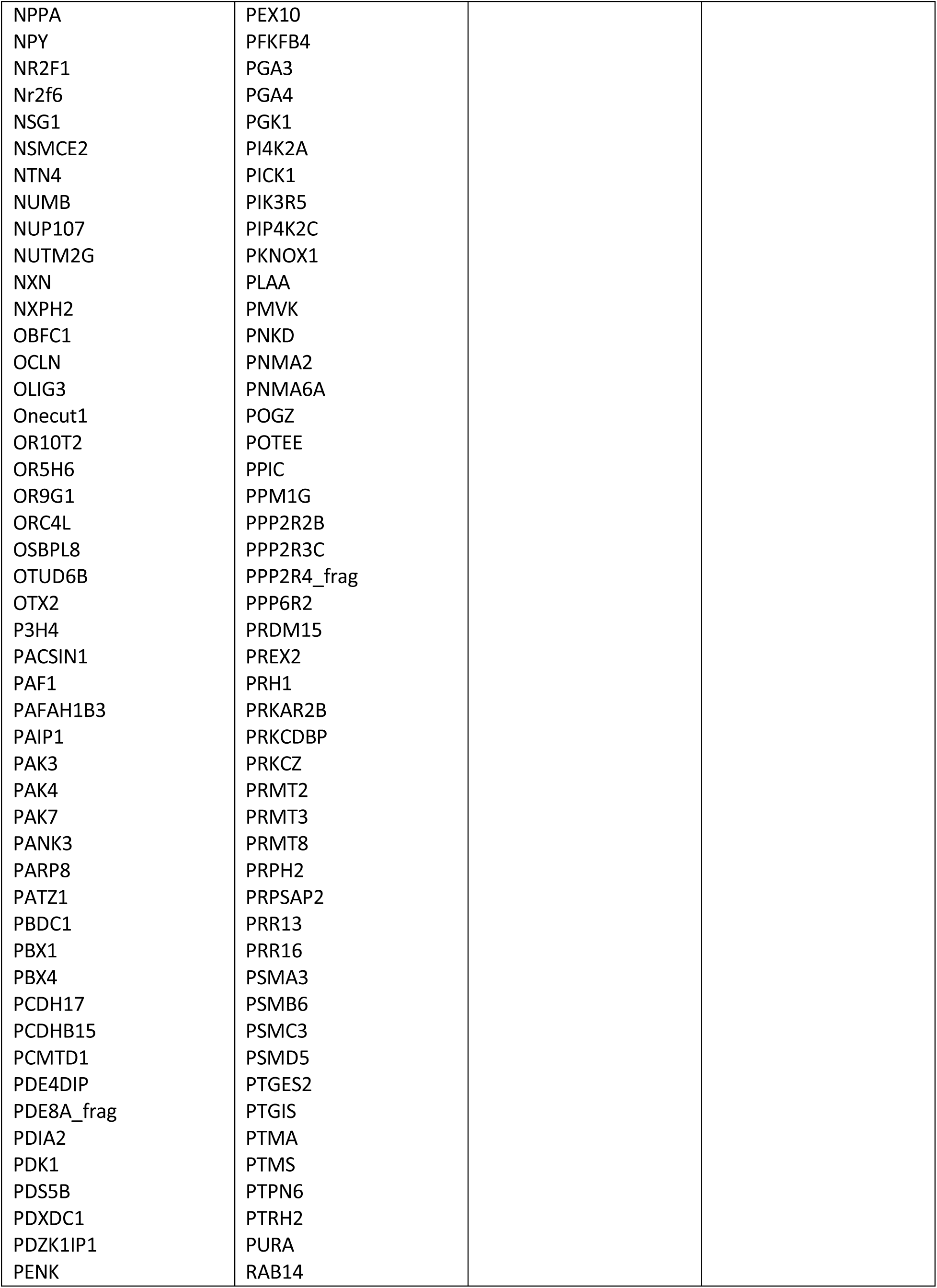
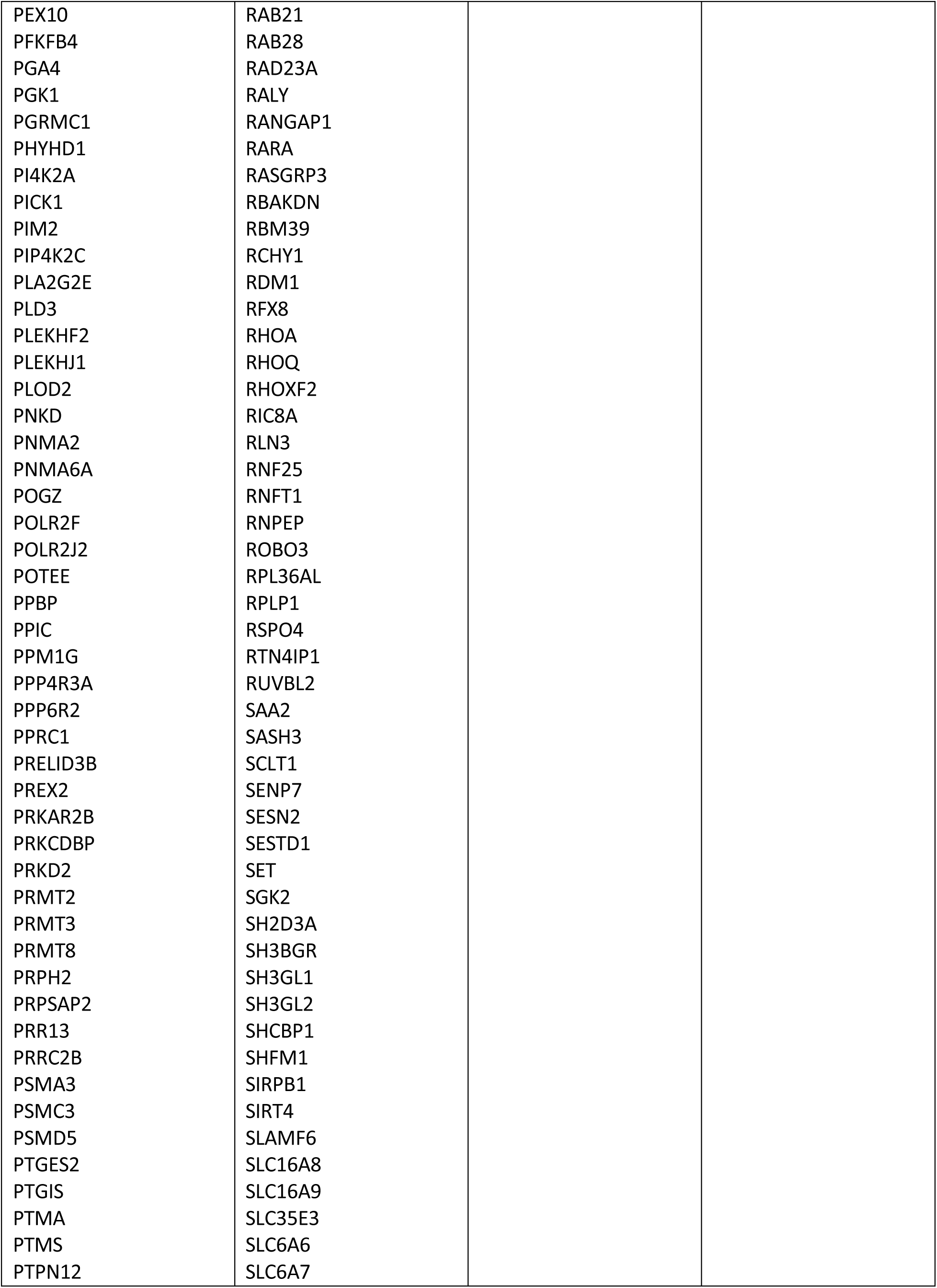
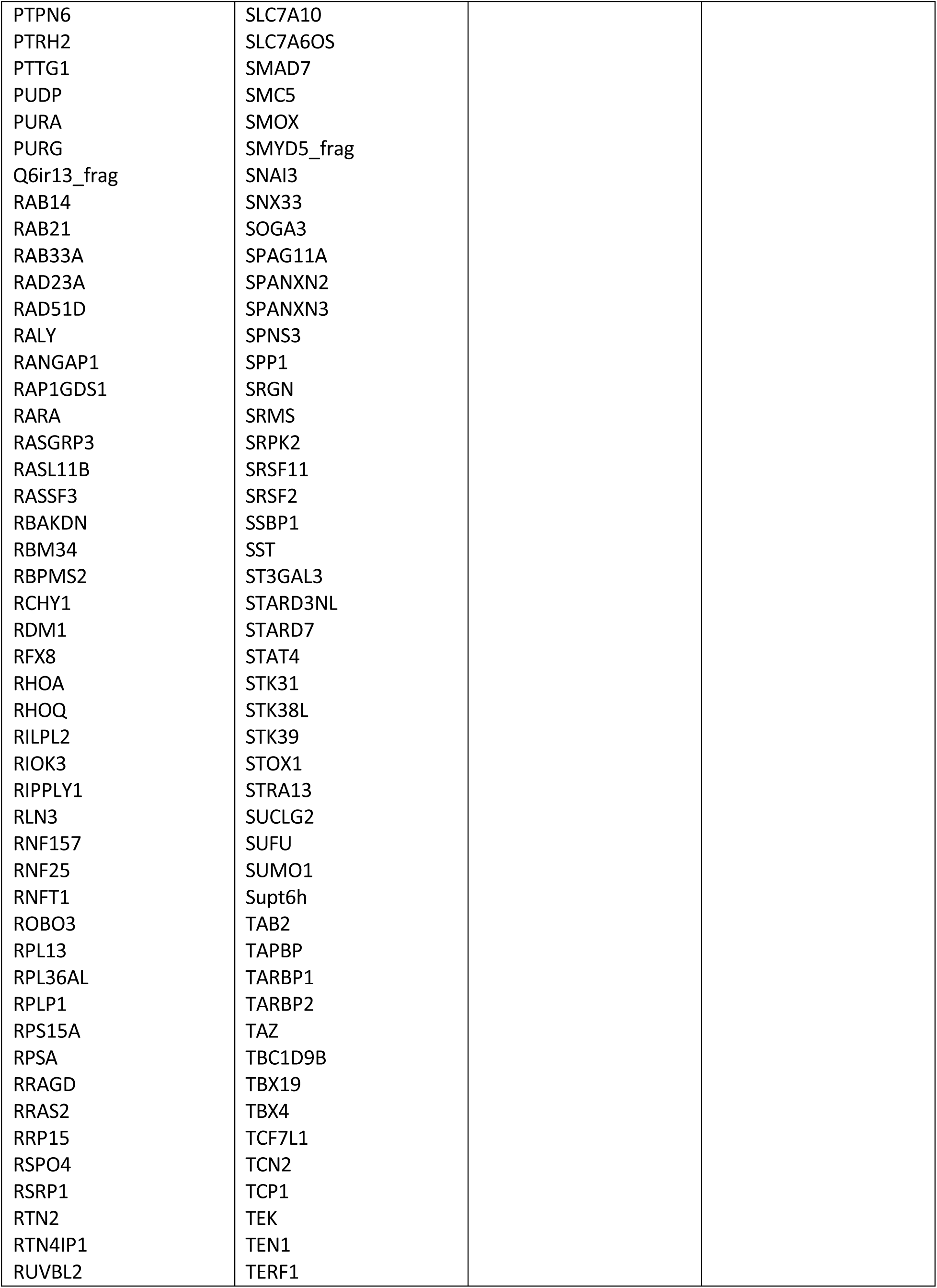
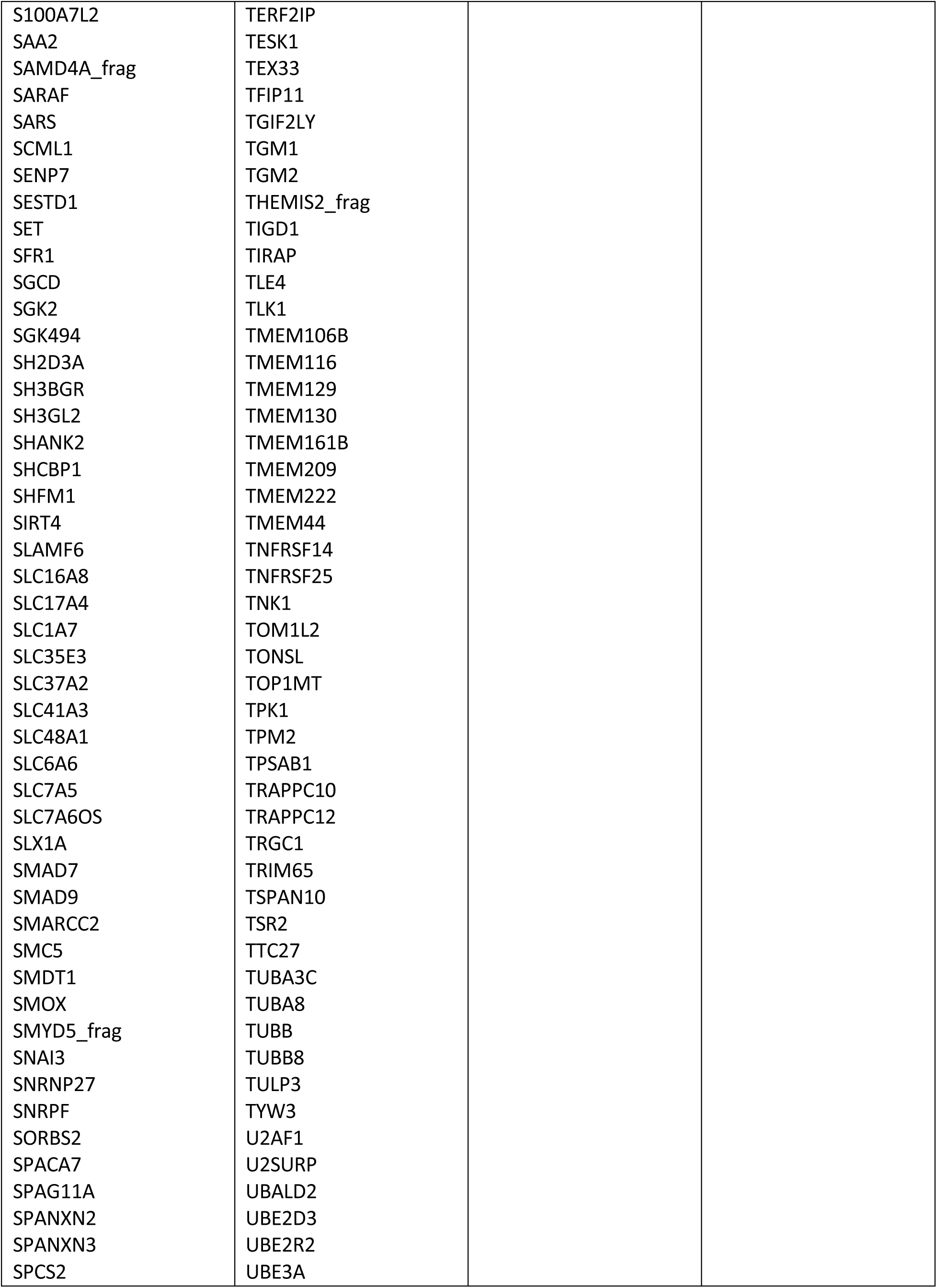
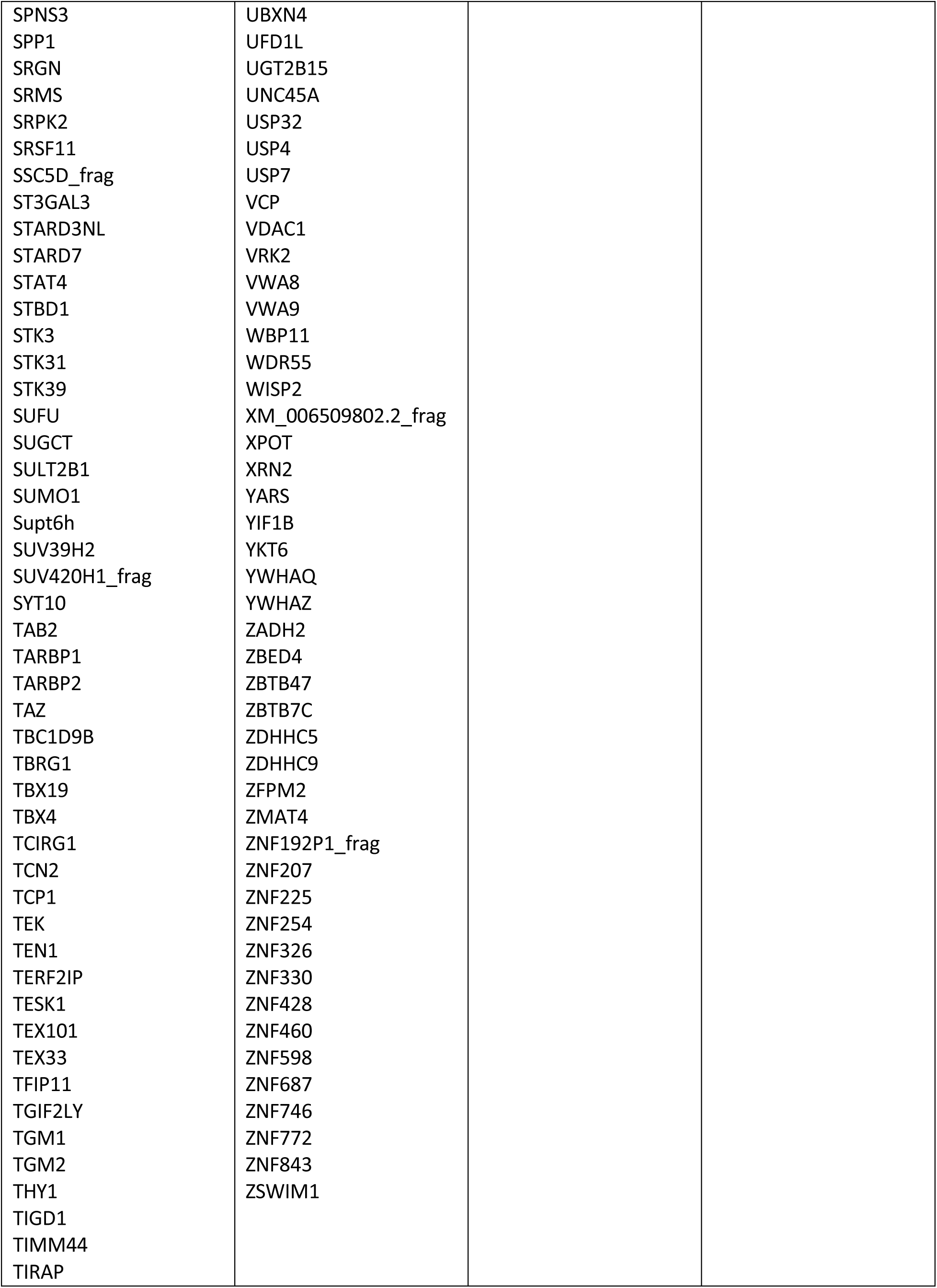
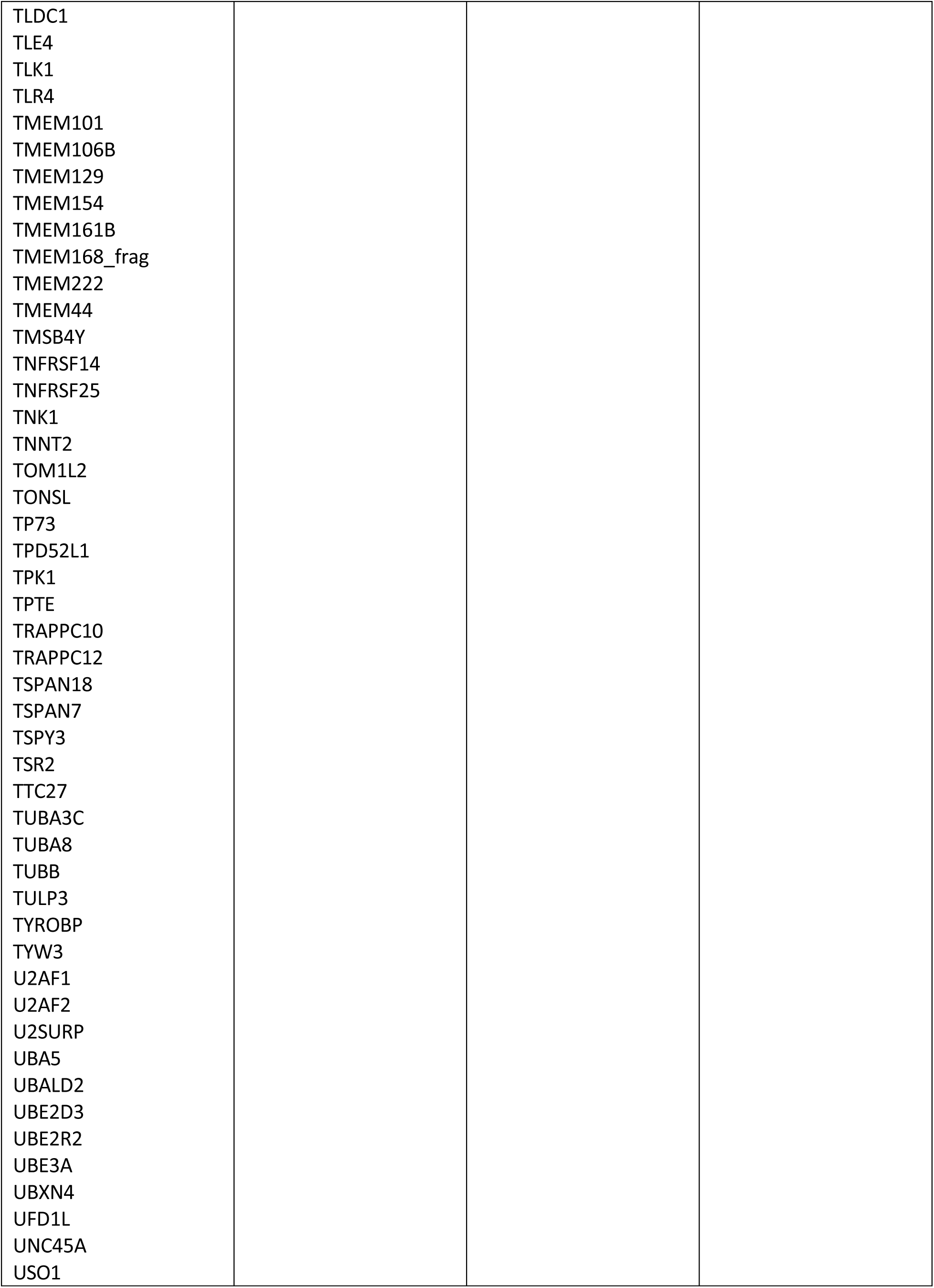
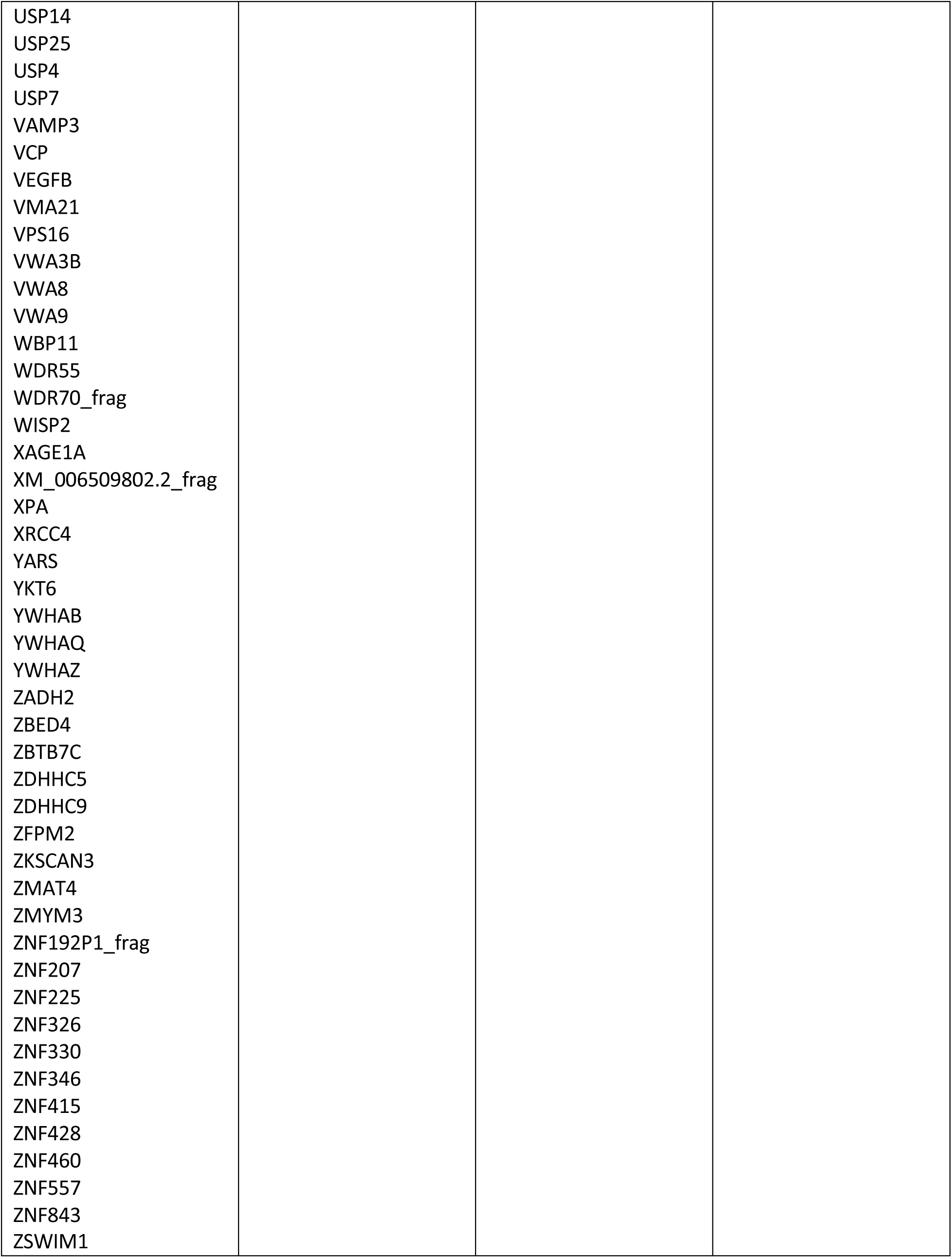
Human protein (autoantigen) sets recognized by both Ig and E. coli-captured Ig from SAH livers

**Table S4.** Cellular components recognized by IgG or IgA antibodies extracted from the diseased liver tissues (Observed vs. Expected)

| Antigen enriched cellular component | Cytosol | Cytoplasm | Nucleus | Mitochondrion | Focal adhesion | Extracellular exosome | Mitochondrial intermembrane space | Ruffle membrane |
| --- | --- | --- | --- | --- | --- | --- | --- | --- |
| SAH (IgG) | P<0.0001 | P<0.0001 | P<0.0001 | P<0.05 | P<0.05 | P<0.05 | P<0.05 | P<0.05 |
| SAH (IgA) | P<0.05 | P<0.05 | ns | ns | ns | ns | ns | ns |
| AC (IgG) | ns | ns | ns | ns | ns | ns | ns | ns |
| AC (IgA) | ns | ns | ns | ns | ns | ns | ns | ns |
| HBV (IgG or IgA) | ns | ns | ns | ns | ns | ns | ns | ns |
| HCV (IgG or IgA) | ns | ns | ns | ns | ns | ns | ns | ns |
| NASH (IgG or IgA) | ns | ns | ns | ns | ns | ns | ns | ns |
| AIH (IgG or IgA) | ns | ns | ns | ns | ns | ns | ns | ns |

**Table S5.** Cellular components recognized by IgM and E. coli enriched IgM extracted from PBC liver tissues.

| Ig from PBC livers |  |  |  |  |
| --- | --- | --- | --- | --- |
| Antigen enriched cellular component | Golgi membrane | Palmitoyltransferase complex | Extracellular exosome | Intrinsic component of Golgi membrane |
| PBC-IgM | P<0.01 | P<0.01 | P<0.05 | P<0.05 |

| E. Coli enriched Ig from PBC livers |  |  |
| --- | --- | --- |
| Antigen enriched cellular component | Cytosol | Protein acetyltransferase complex |
| PBC-IgM | P<0.001 | P<0.05 |

